# SEL1L-HRD1 interaction is prerequisite for the formation of a functional HRD1 ERAD complex

**DOI:** 10.1101/2023.04.13.536796

**Authors:** Liangguang Leo Lin, Xiaoqiong Wei, Huilun Helen Wang, Brent Pederson, Mauricio Torres, You Lu, Zexin Jason Li, Xiaodan Liu, Hancheng Mao, Hui Wang, Zhen Zhao, Shengyi Sun, Ling Qi

## Abstract

The SEL1L-HRD1 protein complex represents the most conserved branch of endoplasmic reticulum (ER)-associated degradation (ERAD); however, definitive evidence for the importance of SEL1L in HRD1 ERAD is lacking. Here we report that attenuation of the interaction between SEL1L and HRD1 impairs HRD1 ERAD function and has pathological consequences in mice. Our data show that *SEL1L* variant *p.Ser658Pro* (*SEL1L*^*S*658*P*^) previously identified in Finnish Hound suffering cerebellar ataxia is a recessive hypomorphic mutation, causing partial embryonic lethality, developmental delay, and early-onset cerebellar ataxia in homozygous mice carrying the bi-allelic variant. Mechanistically, *SEL1L*^*S*658*P*^ variant attenuates the SEL1L-HRD1 interaction and causes HRD1 dysfunction by generating electrostatic repulsion between SEL1L F668 and HRD1 Y30 residues. Proteomic screens of SEL1L and HRD1 interactomes revealed that the SEL1L-HRD1 interaction is prerequisite for the formation of a functional HRD1 ERAD complex, as SEL1L recruits not only the lectins OS9 and ERLEC1, but the E2 UBE2J1 and retrotranslocon DERLIN, to HRD1. These data underscore the pathophysiological importance and disease relevance of the SEL1L-HRD1 complex, and identify a key step in organizing the HRD1 ERAD complex.

## HIGHLIGHTS

1. *SEL1L*^*S*658*P*^ variant is a disease-causing recessive hypomorphic mutation in mice.
2. SEL1L^S658P^ causes HRD1 ERAD defects by attenuating SEL1L-HRD1 interaction.
3. SEL1L^S658P^ generates electrostatic repulsion between F668 SEL1L and Y30 HRD1.
4. SEL1L-HRD1 interaction is prerequisite for a functional HRD1 ERAD complex.
5. Proteomic screens identify SEL1L-dependent and -independent HRD1 interactors.

## INTRODUCTION

In eukaryotes, approximately 30% of all newly synthesized proteins pass through the endoplasmic reticulum (ER), where they undergo folding and maturation. Although a signification subfraction fails to fold properly, evolutionary biology has accounted for this, as many of these proteins are ultimately disposed of by a quality-control process known as ER-associated degradation (ERAD) ^1–3^. While ERAD has been implicated in over 70 human diseases ^4^, our understanding of the significance of the ERAD machinery, as well as its underlying mechanism, in disease pathogenesis remains limited.

Among many different ERAD machineries, the suppressor of lin-12-like (SEL1L, known as Hrd3p in yeast)-HMG-CoA reductase degradation 1 (HRD1) complex represents the most conserved branch of ERAD from yeast to humans ^5–8^. SEL1L acts as the obligatory cofactor for the E3 ligase HRD1. In both yeast and mammals, Hrd3p/SEL1L not only regulates the stability of Hrd1p/HRD1 ^5, 9, 10^, but also provides a scaffold for other ERAD components, including lectins Os9p/OS9 (Osteosarcoma amplified 9) and ERLEC1 (ER lectin 1), cofactors Usa1/HERP1 (U1 SNP1-associated protein 1/Homocysteine-induced ER protein 1) and FAM8A1 (Family with sequence similarity 8 member A1), retrotranslocon Der1p/DERLIN (Degradation in the endoplasmic reticulum 1), E2 Ubc6p/UBE2J1 (Ubiquitin-conjugating enzyme E2 J2), a cytosolic E2 UBC2 (UBE2G2 in mammals), and ATPase CDC48p/VCP (Valosin containing protein) ^10–24^. Biochemical and genetic studies in yeast and mammalian cells have shown that misfolded proteins in the ER are recruited to Hrd1p/HRD1 through Yos9/Os9 and Hrd3p/SEL1L ^25^, retrotranslocated and ubiquitinated through the activities of Hrd1p-Der1p retrotranslocon and the E2 Ubc6p/UBE2J1 ^19, 24, 26^ for proteasomal degradation. How this HRD1 protein complex is assembled together remains vague.

In mammals, global or acute deletion of *Sel1L* or *Hrd1* in germline and adult mice leads to embryonic or premature lethality, highlighting the crucial roles of SEL1L and HRD1 in both embryonic development and adult stages ^9, 27–29^. Recent studies using cell type-specific mouse models have further demonstrated the vital importance of SEL1L and HRD1 in a cell type- and substrate-specific manner in physiological process ^2, 30–50^. However, questions about the role and importance of SEL1L in HRD1 ERAD are still lingering. Indeed, some recent findings have questioned the necessity of SEL1L: (1) Hrd3p/SEL1L function can be bypassed by overexpressing Hrd1p/HRD1 in both yeast and mammalian cell culture systems ^10, 32, 51, 52^; (2) In yeast, Hrd3p is dispensable for Hrd1-substrate, Hrd1-E2 interaction, although it is required for the E3 ubiquitination activity of Hrd1 (or substrate ubiquitination) ^18^; (3) in mammals, additional SEL1L-independent HRD1 complexes have been proposed or identified, such as the HRD1-FAM8A1 complex ^23, 53^. Hence, although there has been ample evidence for the importance of individual SEL1L and HRD1 proteins, definitive evidence for the importance of this protein complex remains lacking in mammals.

In analyzing disease causality of *SEL1L*^*S*658*P*^ variant (*p.Ser658Pro;* c.1972T>C, NM_005065), first reported in 2012 in Finnish Hound suffering cerebellar ataxia ^54^, we serendipitously uncovered a prerequisite step in the formation of a functional HRD1 ERAD complex. We showed that this SEL1L variant is disease causal in mice due to a partial loss of function of, i.e. hypomorphic, ERAD. Mechanistically, *SEL1L*^*S*658*P*^ causes HRD1 dysfunction by attenuating SEL1L-HRD1 interaction via F668 SEL1L-Y30 HRD1. Our unbiased proteomics screens of SEL1L and HRD1 interactomes revealed that SEL1L is required for the formation of an HRD1 functional complex as it recruits lectins OS9 and ERLEC1, the E2 enzyme UBE2J1 and retrotranslocon DERLIN to HRD1 for substrate recruitment, ubiquitination and retrotranslocation, respectively. Together, these biochemical and pathological data demonstrate the importance of SEL1L in HRD1 ERAD and its underlying mechanism.

## RESULTS

### Generation of knock-in (KI) mice carrying *SEL1L*^*S*658*P*^variant

To establish disease causality of the *SEL1L*^*S*658*P*^ variant, we generated knock-in (KI) mice carrying this variant on the C57BL/6J/SJL mix genetic background using the CRISPR/Cas9-based methods (Figure S1A). Two independent founders were established by PCR and sequencing (Figure S1B and S1C) and then used to generate bi-allelic homozygous KI mice by intercrossing the F1 generations of a cross between the founders and C57BL/6J WT mice. Each founder line was characterized independently. As similar results were obtained from each founder line, the results were combined.

Much to our surprise, we obtained homozygous KI pups at a frequency of less than 11% in 247 pups of over 30 litters from 12 breeding pairs, less than half of the expected Mendelian transmission of a recessive trait (25%) (Figure S1D). Timed pregnancy followed by PCR-genotyping of the embryos at embryonic day (E) 10.5-12.5 during organogenesis and at E14.5-16.5 during body-mass growth and organ maturation showed that empty deciduae (i.e. reabsorbed embryos) were found at E10.5 and all turned out to be KI embryos (Figure 1A). Indeed, the frequency of KI embryos in a total of over 100 embryos was progressively reduced from 17% at E10.5-12.5 to 14% at E14.5-16.5, while the frequency of heterozygous mice (HET) maintained at ∼58% (Figure S1D). Hence, these data demonstrate that the *SEL1L*^*S*658*P*^ allele is a recessive mutation and causes partial embryonic lethality at ∼50% frequency in this mixed genetic background.

**Figure 1.**
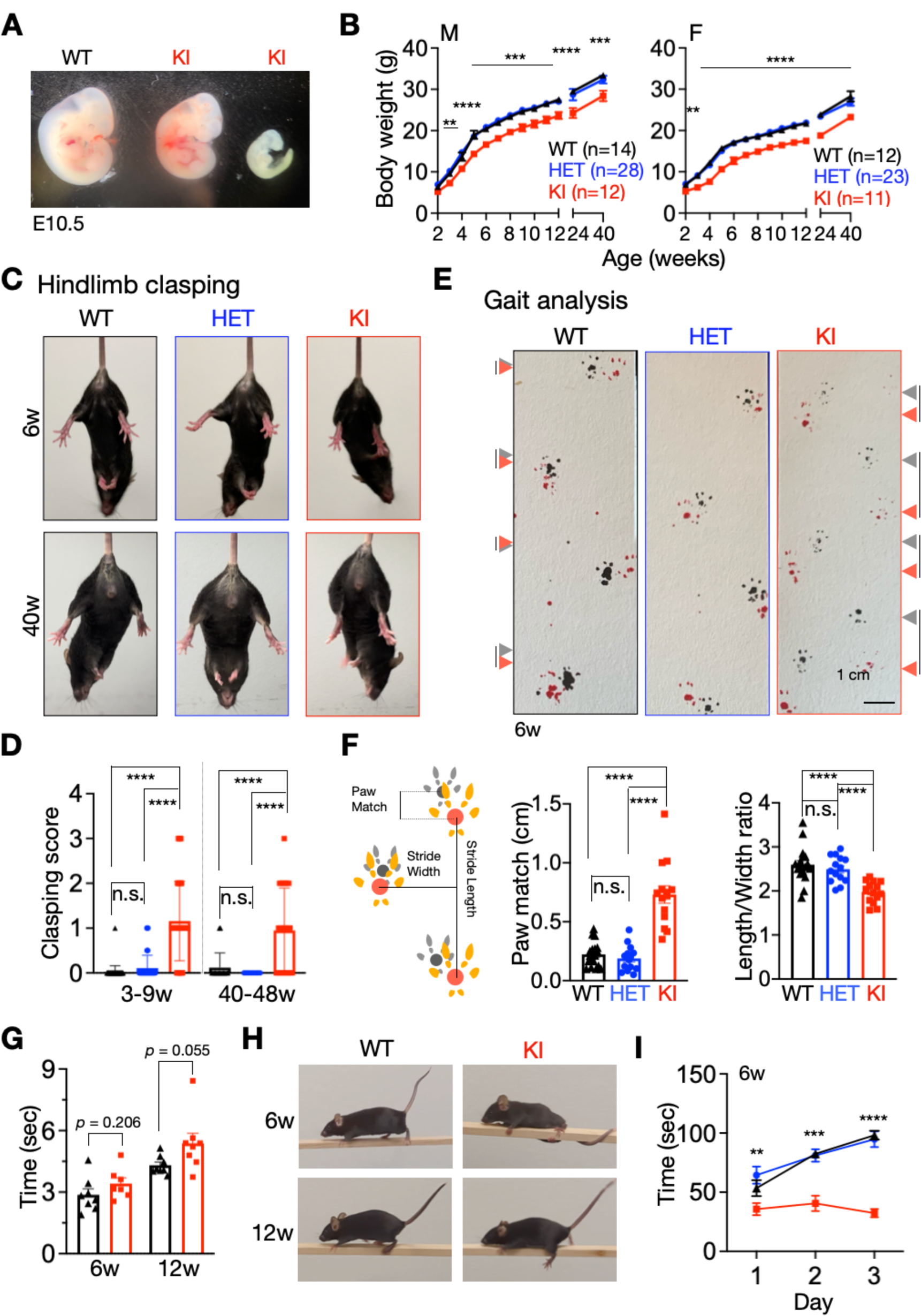
*SEL1L*^*S*658*P*^ KI mice exhibit partial embryonic lethality, developmental delay and early-onset, non-progressive, ataxia. **(A)** Gross appearance of *SEL1L*^*S*658*P*^KI embryos (one dead, right) compared with their WT littermates at E10.5. **(B)** Growth curve. n = 12-28 mice for male and n = 11-23 mice for female per group. **(C-D)** Representative pictures of hindlimb clasping (C) and score quantitation (D) of 3-9 (n = 49, 51 and 38 for WT, HET and KI) and 40-48 (n = 17, 8 and 20 mice for WT, HET and KI)-week-old littermates of both genders. **(E-F)** Representative pictures (E), cartoon schematic of paw prints (left, F) and quantitation of gait analysis (right, F) of 6-week-old littermates of both genders. The grey and red arrows in (E) indicate forelimb and hindlimb, respectively; and the lines between grey and red arrow indicate the distance between the two limbs (n = 14-18 mice per group). **(G)** Quantitation of balance beam test from mice at 6 and 12 weeks of age (n = 7-8 per group). **(H)** Representative pictures of slips on balance beam of 6 and 12-week-old KI mice. **(I)** Quantitation of rotarod test from 6-week-old mice with 3 days training (n = 7, 12 and 6 mice for WT, HET and KI). Values, mean ± SEM. n.s., not significant; **p<0.01, ***p<0.001 and ****p<0.0001 by two-way ANOVA followed by Tukey’s multiple comparisons test (B, I) and one-way ANOVA followed by Tukey’s post hoc test (D, F, G).

### Mild developmental delay of *SEL1L*^*S*658*P*^ KI mice

In the surviving cohorts, KI mice exhibited modest growth retardation by ∼10-15% in both sexes, starting at 3-4 weeks through the first 40 weeks of age (Figure 1B). At 5 weeks of age, the KI mice of both sexes were slightly shorter in body length compared to that of WT littermates (Figure S2A). This growth delay was not due to reduced food intake or associated with abnormal blood glucose levels, as they were comparable among the cohorts (Figures S2B and S2C). Much to our surprise, blind histological examination of peripheral tissues of 5-week-old KI mice by a pathologist revealed no obvious histological abnormalities in the pancreas, liver, kidneys, inguinal white adipose tissue (iWAT), and brown adipose tissue (BAT) (Figures S2D, S2E and S2G). Biochemical analyses of serum revealed no significant damage in the livers of both sexes based on the levels of alanine transaminase (ALT), alkaline phosphatase (ALP) and total bilirubin (TBIL) (Figure S2F), or in the kidneys based on blood urea nitrogen (BUN) and creatinine levels (CREA) (Figure S2H). Of note, although ALT level was significantly increased in male KI mice compared to that of WT littermates, it was still within the normal range (24.3-115.3 U/L) (Figure S2F). Hence, these data demonstrate that *SEL1L*^*S*658*P*^ is a recessive hypomorphic mutation causing mild growth retardation, while having no major effect on peripheral tissues in mice.

### Early-onset, non-progressive, ataxia in *SEL1L*^*S*658*P*^ KI mice

KI mice developed abnormal hind limb-clasping reflexes when suspended by their tails, starting at 3-9 weeks of age (Figures 1C and 1D). Intriguingly, the phenotype remained mild for up to 48 weeks of age, pointing to the non-progressive nature of the disease (Figures 1C and 1D). In addition, unlike WT mice showing a normal symmetric gait characterized by the overlapping of hindlimb and forelimb paw prints, KI mice exhibited asymmetric gait patterns characterized by the lagging of the hindlimbs and reduction in stride length at 6 weeks of age (arrows, Figures 1E and 1F). Interestingly, while the balance beam test showed that KI mice at 6 and 12 weeks of age had no problems completing the test compared to WT littermates (Figures 1G), some KI mice slipped and lost their balance while crossing the beam (Figure 1H and Video S1), pointing to an impairment of balance. Moreover, KI mice showed worse performance of motor coordination in the rotarod tests compared to WT littermates even with training at 6 weeks of age (Figures 1I and S2I). HET mice were comparable to WT littermates in all above tests (Figure 1 and not shown), in line with the autosomal recessive nature of the variant. These data show that *SEL1L*^*S*658*P*^ KI mice exhibit growth retardation, and signs of early onset non-progressive ataxia, establishing the disease-causality and pathogenicity of this allele.

### Microcephaly and a mild reduction of Purkinje cells in *SEL1L*^*S*658*P*^ KI mice

We next further explored the molecular changes in the brain that may underlie the pathogenesis of early-onset ataxia. At 5 weeks of age, KI mice had reduced brain weight by ∼10 % compared to those of age-matched WT and HET littermates (Figure 2A). MRI analyses revealed smaller cerebellum and cortex in KI mice by ∼10% relative to those of WT littermates, i.e. microcephaly (Figure 2B and S3A). Blind histological examination by a pathologist revealed no obvious histological abnormalities in various brain regions, including the cortex, of KI mice at 5-weeks of age (Figures S3B and S3C). Immunostaining of markers for neurons (NeuN), astrocytes (GFAP), and microglia (IbaI) also showed comparable staining pattens in the whole brain of KI vs. WT mice at 5 weeks of age (Figures S3D and S3E), excluding the possibility of massive cell death and neuroinflammation. Hence, these data show that KI mice exhibit microcephaly, with no massive cell death or inflammation in the brain.

**Figure 2.**
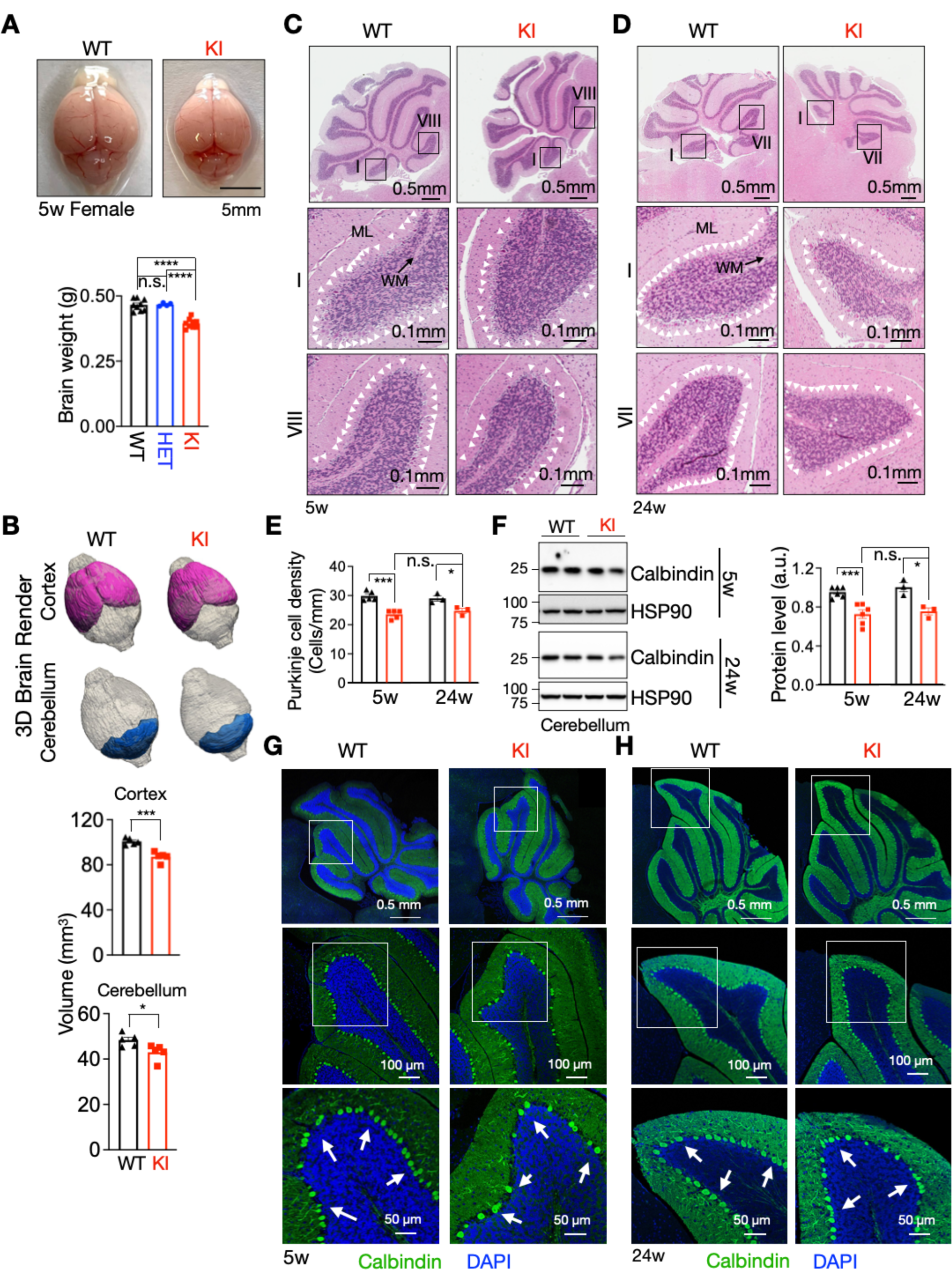
*SEL1L*^*S*658*P*^ KI mice exhibit microcephaly and a mild reduction of Purkinje cells. **(A)** Representative pictures of brains of 5-week-old mice, with quantitation of brain weight shown below (n = 9, 4 and 9 mice for WT, HET and KI). **(B)** 3D rendering of cortex and cerebellum from 5-week-old mice, with quantitation shown below (n = 5 mice per group). **(C-E)** Hematoxylin & eosin (H&E) stained sagittal sections of the cerebellum from 5-(C) and 24-(D) week-old mice, with quantitation of Purkinje cell density shown in (E) (n = 3-5 mice per group). White arrowheads, Purkinje cells. ML, molecular layer; MW, white matter. **(F)** Western blot analysis of Calbindin in the cerebellum of 5-and 24-week-old mice with quantitation shown on the right (n = 3-6 mice per group). **(G-H)** Representative confocal images of Calbindin (green) in the cerebellum of 5-(G) and 24-(H) week-old mice. White arrows, Purkinje cells (n = 3 mice per group). Values, mean ± SEM. n.s., not significant; *p<0.05 and ***p<0.001 by one-way ANOVA followed by Tukey’s post hoc test (A) and two-tailed Student’s *t*-test (B, E, F).

Given the ataxia phenotype of the KI mice (Figure 1), we next examined the cerebellar region consisted of Purkinje neurons as alterations in Purkinje cells have been directly associated with cerebellar ataxia^55–57^. Purkinje cells are arranged in a single layer in the cerebellar cortex, with their dendrites facing the molecular layer (ML) and their axons facing the white matter (WM) (Figures 2C and 2D). Quantitation of the number of Purkinje cells based on their unique location and morphology in H&E-stained sagittal sections of the cerebellum revealed a mild 15-20% reduction in KI mice at 5 weeks of age (Figures 2C and 2E). This reduction did not worsen with age as a similar level of reduction was observed at 24 weeks of age (Figures 2D and 2E). This result was further confirmed by the quantitation of the Purkinje cell marker Calbindin using Western blot (Figure 2F) and immunostaining at both 5 and 24 weeks of age (Figures 2G and 2H). Hence, our data show that *SEL1L*^*S*658*P*^ is associated with microcephaly and a mild, but non-progressive, reduction of Purkinje cells starting at a young age.

### Cellular adaptation in the cerebellum of KI mice

As ER stress was implicated in the pathogenesis of Finnish Hounds carrying biallelic *SEL1L*^*S*658*P*^ variant in the earlier study ^54^, we next asked whether and how ER homeostasis was affected in the cerebellum of *SEL1L*^*S*658*P*^ KI mice. In line with our previous study that UPR sensor IRE1α is an ERAD substrate ^32^, its protein level was increased by ∼4 folds; however, neither IRE1α phosphorylation nor splicing of its downstream effector *Xbp1* mRNA was elevated in the cerebellum of 5-week-old KI mice (Figure 3A). While phosphorylation of another UPR sensor PERK and its downstream effector was enhanced by 1.5-2 folds, quantitation of the ration of phosphorylated to total proteins showed a subtle activation of the PERK pathway (Figure 6B).

**Figure 3.**
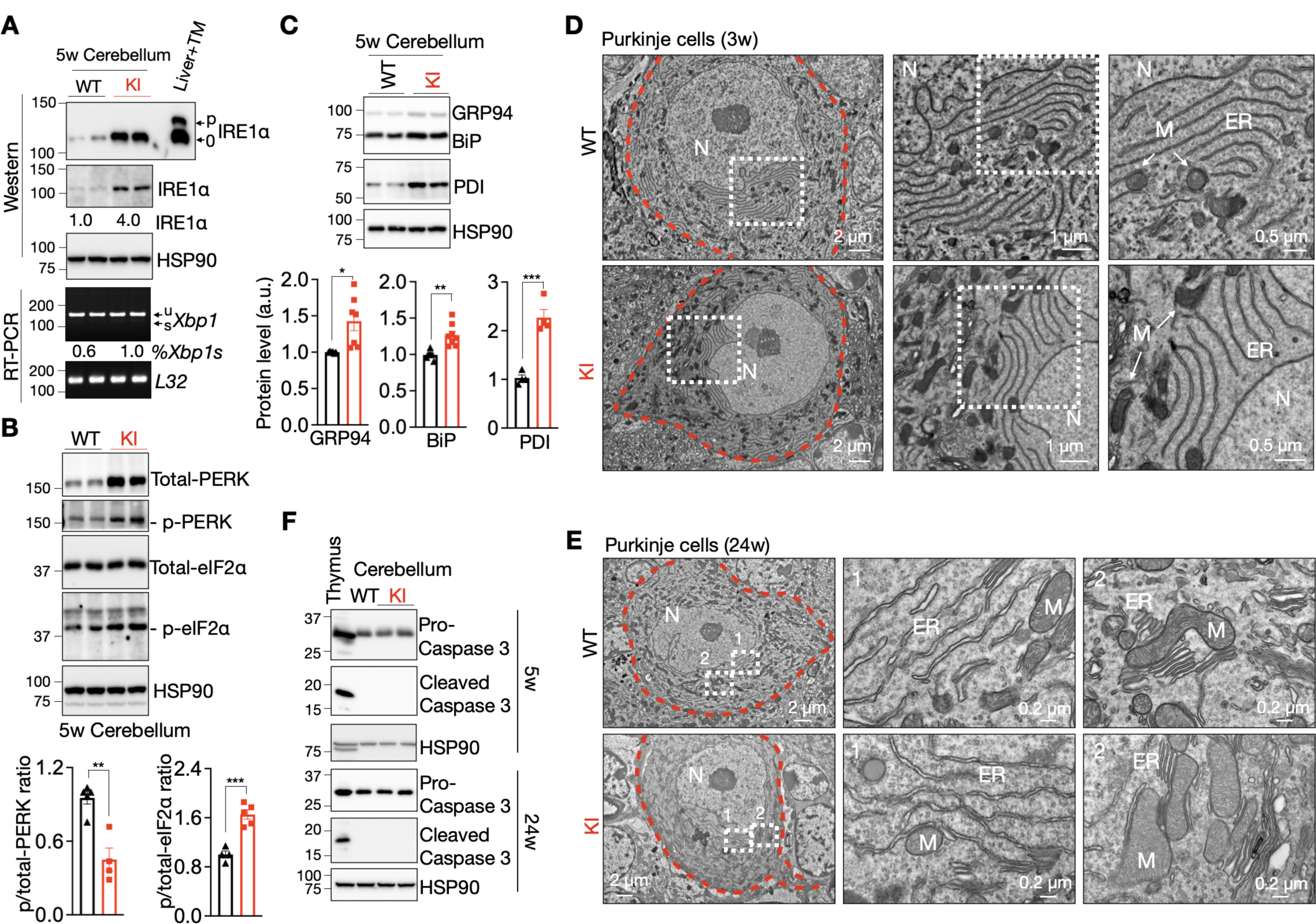
Subtle alteration of ER homeostasis in the cerebellum of *SEL1L*^*S*658*P*^ KI mice. **(A)** Western blot analysis of IRE1α phosphorylation (top) and RT-PCR of *Xbp1* splicing in the cerebellum of 5-week-old mice. 0/p, non-/phosphorylation; u/s, un-/spliced *Xbp1*. Quantitation of IRE1α and the ratio of spliced to total *Xbp1* shown below the gel as mean (n = 4-5 mice per group). **(B)** Western blot analysis of PERK and eIF2α phosphorylation in the cerebellum of 5-week-old mice with quantitation shown below (n = 4-5 mice per group). **(C)** Western blot analysis of ER chaperones GRP94, BiP and PDI in the cerebellum of 5-week-old mice with quantitation shown (n = 4-8 mice per group). **(D-E)** Representative TEM images of Purkinje cells (outlined by red dotted lines) of 3-(D) and 24-(E) week-old mice. N, nucleus; M, mitochondria. n = 2 mice each genotype at each age. **(F)** Western blot analysis of cell death marker Caspase 3 in the cerebellum of 5- and 24-week-old mice. Thymus, a positive control (n = 3 mice each genotype). Values, mean ± SEM. *p<0.05, **p<0.01 and ***p<0.001 by two-tailed Student’s *t*-test (B, C).

Moreover, protein levels of key ER chaperones such as BiP, GRP94 and protein disulfide isomerase (PDI) were modestly increased by 20-40 and 100%, respectively, in the cerebellum of KI mice relative to those in WT littermates (Figure 3C). In keeping with the subtle changes of UPR, transmission electron microscopic (TEM) examination of Purkinje cells revealed largely normal ER sheet morphology in both 3- and 24-week-old KI mice (Figures 3D and 3E). Cleaved caspase 3, a marker of apoptosis, was undetectable in the cerebellum of KI mice at either 5 or 24 weeks of age (Figure 3F). Taken together, these data demonstrate that neurons including Purkinje cells can adapt to the expression of *SEL1L*^*S*658*P*^ without eliciting an overt ER stress or cell death.

### *SEL1L*^*S*658*P*^ is a hypomorphic variant with impaired ERAD function

We next asked whether *SEL1L*^*S*658*P*^ causes HRD1 ERAD dysfunction. We first tested the turnover and aggregation of a known ERAD substrate, a disease mutant of pro-arginine vasopressin (proAVP) at residue 57 (Gly-to-Ser, Gly57Ser) ^35, 37^. In line with our previous studies ^35, 37^, proAVP Gly57Ser protein accumulated and formed HMW aggregates in *SEL1L^-/-^* HEK293T cells (Figure 4A, lane 2 vs. 1), which was attenuated upon the over-expression of SEL1L WT (lane 3-5 vs. 2), but not SEL1L S658P (lane 6-8 vs. 3-5, Figure 4A and quantitated in Figure 4B).

**Figure 4.**
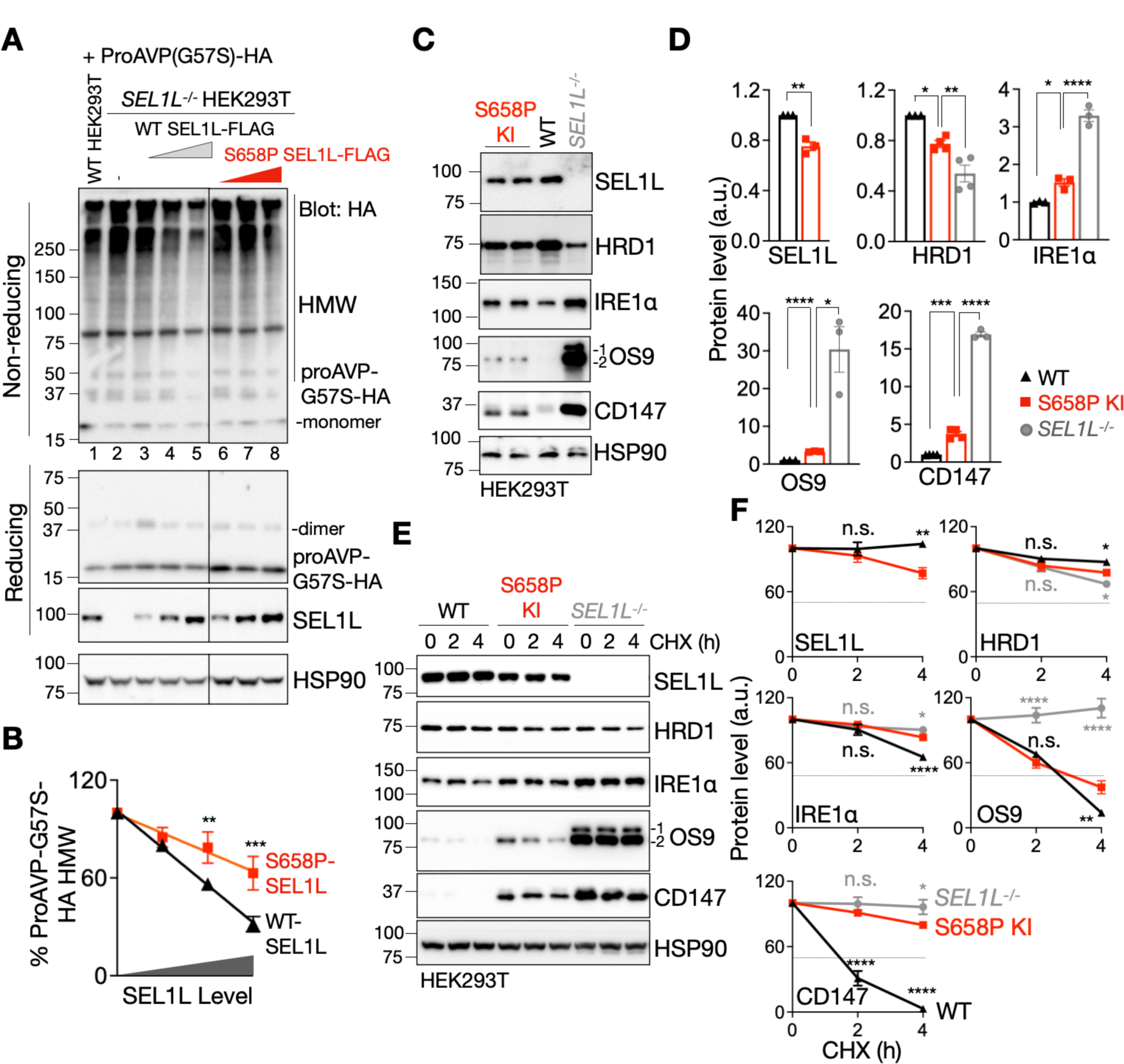
*SEL1L*^*S*658*P*^ is a hypomorphic variant with impaired ERAD function. **(A-B)** Reducing and non-reducing SDS-PAGE and Western blot analyses of proAVP-G57S high molecular-weight (HMW) aggregates in WT or *SEL1L^-/-^* HEK293T cells transfected with indicated SEL1L-FLAG constructs in a dose-dependent manner, with quantitation of proAVP-G57S-HA HMW shown in (B) (n = 7 and 4 independent samples for WT and S658P SEL1L-FLAG). Two panels in (A) were from the same experiment with the irrelevant lanes in the middle cut off. **(C-D)** Western blot analysis of SEL1L, HRD1 and known endogenous substrates IRE1α, OS9 and CD147 in WT, *SEL1L*^*S*658*P*^ KI or *SEL1L^-/-^* HEK293T cells, with quantitation shown in (D) (n = 3-4 independent samples each genotype). **(E-F)** Cycloheximide (CHX) chase analysis of SEL1L, HRD1 and known endogenous substrates IRE1α, OS9 and CD147 in WT, *SEL1L*^*S*658*P*^ KI or *SEL1L^-/-^* HEK293T cells, with quantitation shown in (F) (n = 3-5 independent samples each genotype). Values, mean ± SEM. n.s., not significant; *p<0.05, **p< 0.01, ***p<0.001 and ****p<0.0001 by two-way ANOVA followed by Dunnett’s multiple comparisons test (B, F) and one-way ANOVA followed by Dunnett’s multiple comparisons test (D).

We then generated KI HEK293T cells carrying the bi-allelic *SEL1L*^*S*658*P*^ variant (Figures S4A and S4B) and compared them to WT and *SEL1L^-/-^* HEK293T cells. Both KI and KO cell lines were generated using CRISPR/Cas9 system. Protein levels of SEL1L and HRD1 were reduced by 20 and 30%, respectively, in *SEL1L*^*S*658*P*^ KI HEK293T cells compared to WT HEK293T cells (Figures 4C and 4D). Indeed, *SEL1L*^*S*658*P*^ attenuated the stability of both SEL1L and HRD1 proteins (Figures 4E and 4F). Moreover, *SEL1L*^*S*658*P*^ expression led to a significant accumulation, as a result of protein stabilization, of several known ERAD substrates such as IRE1α, OS9, and CD147 in KI cells compared to those in WT HEK293T cells, but to a much less extent when compared to those in *SEL1L^-/-^* HEK293T cells (Figures 4C-4D and 4E-4F). Hence, these data demonstrate that *SEL1L*^*S*658*P*^ causes partial HRD1 ERAD dysfunction and hence is a hypomorphic variant.

### Sequence and structural analyses of *SEL1L*^*S*658*P*^ variant

We next asked mechanistically how *SEL1L*^*S*658*P*^ affects HRD1 ERAD function. The mutation SEL1L S658P is located at the Sel1-like repeat-C (SLR-C) domain, is highly conserved from Drosophila to humans, but not in yeast (Figures 5A and S5A). Position-specific scoring matrix (PSSM) scores for residue 658 were positive for Ser (green) but negative for Pro (red, Figure S5B), suggesting that Ser has been evolutionarily selected and that Pro at this position might have detrimental effects on SEL1L function. To gain insights into the structural consequences of this variant, we performed the AI-based structural prediction AlphaFold analysis for the human SEL1L (1-722 aa)-HRD1 (1-334 aa)-DERLIN-OS9 protein complex (Figure 5B), based on the Cryo-EM structure of the yeast protein complex (Figure S5C). The structure for the human complex was quite similar to that of the yeast (Figure S5C). Notably, S658 is located on an α-helix in a close proximity to the amphipathic helix of SEL1L ^19, 58^, which directly interacts with the transmembrane 1 (TM1)–TM2 loop of HRD1 (Figures 5B and 5C).

**Figure 5.**
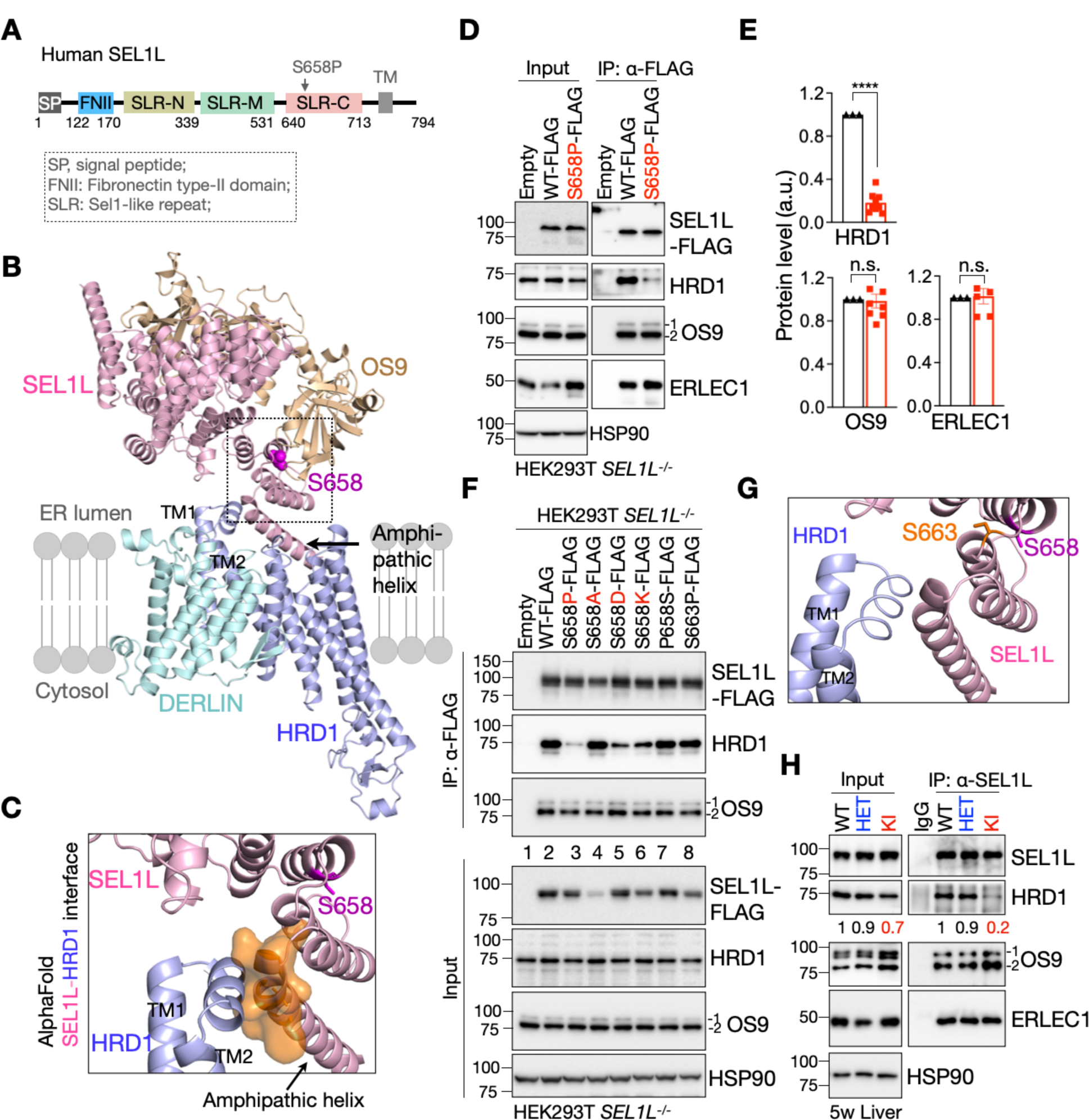
SEL1L^S658P^ attenuates its interaction with HRD1. **(A)** Schematic diagram of human SEL1L with the location of the variant indicated. SP, signal peptide; FNII, fibronectin type II domain; SLR-N/M/C, Sel1-like repeats at N-, Middle- and C-terminal; TM, transmembrane. **(B)** Structural prediction of human SEL1L/OS9/HRD1/DERLIN ERAD complex using AlphaFold2 with SEL1L S658 shown in pink atoms. The arrow indicates the amphipathic helix which interacts with HRD1. **(C)** Side view of a space-filling model of the HRD1-SEL1L interface containing TM1-2 of HRD1 and the amphipathic helix of SEL1L. **(D-E)** Immunoprecipitation of FLAG-agarose in *SEL1L^-/-^* HEK293T cells transfected with indicated SEL1L-FLAG constructs to examine the interaction with HRD1 and OS9/ERLEC1, with quantitation shown in (E) (n = 4-9 independent samples per genotype). **(F)** Immunoprecipitation of FLAG-agarose in *SEL1L^-/-^* HEK293T cells transfected with indicated SEL1L-FLAG constructs to examine the interactions with HRD1 and OS9 (two independent repeats). **(G)** Side view of the HRD1-SEL1L interface. S658 and S663 are indicated in pink and orange, respectively. **(H)** Immunoprecipitation of SEL1L in the livers from 5-week-old WT and KI mice to examine the endogenous SEL1L interaction with HRD1, with quantitation of HRD1 shown below the gel as mean (two independent repeats). Values, mean ± SEM. n.s., not significant; ****p<0.0001 by two-tailed Student’s *t*-test (E).

### *SEL1L*^*S*658*P*^ variant attenuates SEL1L-HRD1 interaction

Based on this modeling, we hypothesized that *SEL1L*^*S*658*P*^ may interfere with the formation of SEL1L-HRD1 protein complex (Figure 5B). In *SEL1L^-/-^*HEK293T cells transfected with FLAG-tagged WT or S658P SEL1L, SEL1L S658P markedly attenuated the interaction between SEL1L and endogenous HRD1 compared to SEL1L WT, while having no effect on its interaction with other components of the complex including OS9 and ERLEC1 (Figures 5D and 5E). Similar results were obtained in *SEL1L^-/-^* HEK293T cells overexpressing both SEL1L-FLAG and HRD1-Myc (Figure S5D). Furthermore, mutation of Ser658 to Ala (S658A) had no effect on the SEL1L-HRD1 interaction (lane 4 vs. 2-3), while S658D or K attenuated the interaction, albeit not as dramatically as S658P (lane 5-6 vs. 2-3, Figure 5F). None of these mutations at SEL1L S658 affected its interaction with OS9 (Figure 5F). Mutating P658 back to S (P658S) rescued the interaction between SEL1L and HRD1 (lane 7, Figure 5F), excluding the possibility of additional mutations outside of S658P. In addition, mutation of a nearby Ser at position 663, 5 amino acids downstream of S658 located at the loop between the two α-helices (Figure 5G), to Pro (S663P) had no apparent effect on SEL1L-HRD1 interaction (lane 8, Figure 5F). In line with the overexpression experiments, SEL1L interaction with HRD1 was reduced by 5 folds in KI livers, while having no effect on the interactions with OS9 and ERLEC1, compared to WT and HET livers (Figure 5H). Hence, we conclude that *SEL1L*^*S*658*P*^ attenuates SEL1L-HRD1 interaction while having no effect on the interaction between SEL1L and lectins.

### *SEL1L*^*S*658*P*^ attenuates the interaction with HRD1 via SEL1L F668 - HRD1 Y30

We then investigated mechanistically how SEL1L S658P attenuates SEL1L-HRD1 interaction. Structral predication revealed a potential physical interaction between SEL1L-F668 (on a neighboring helix) and HRD1-Y30 (on TM1 of HRD1) through an aromatic-aromatic interaction (Figure 6A) in an “edge-to-face” attractive orientation ^59^. Indeed, both F668 and Y30 residues are located within the SEL1L-HRD1 interface (Figure S6A). However, in SEL1L^S658P^, F668-Y30 interaction changed to a “face-to-face” orientation (Figure 6A), which may result in an unfavorable electrostatic repulsion force between the two planar faces of aromatic rings ^59^.

**Figure 6.**
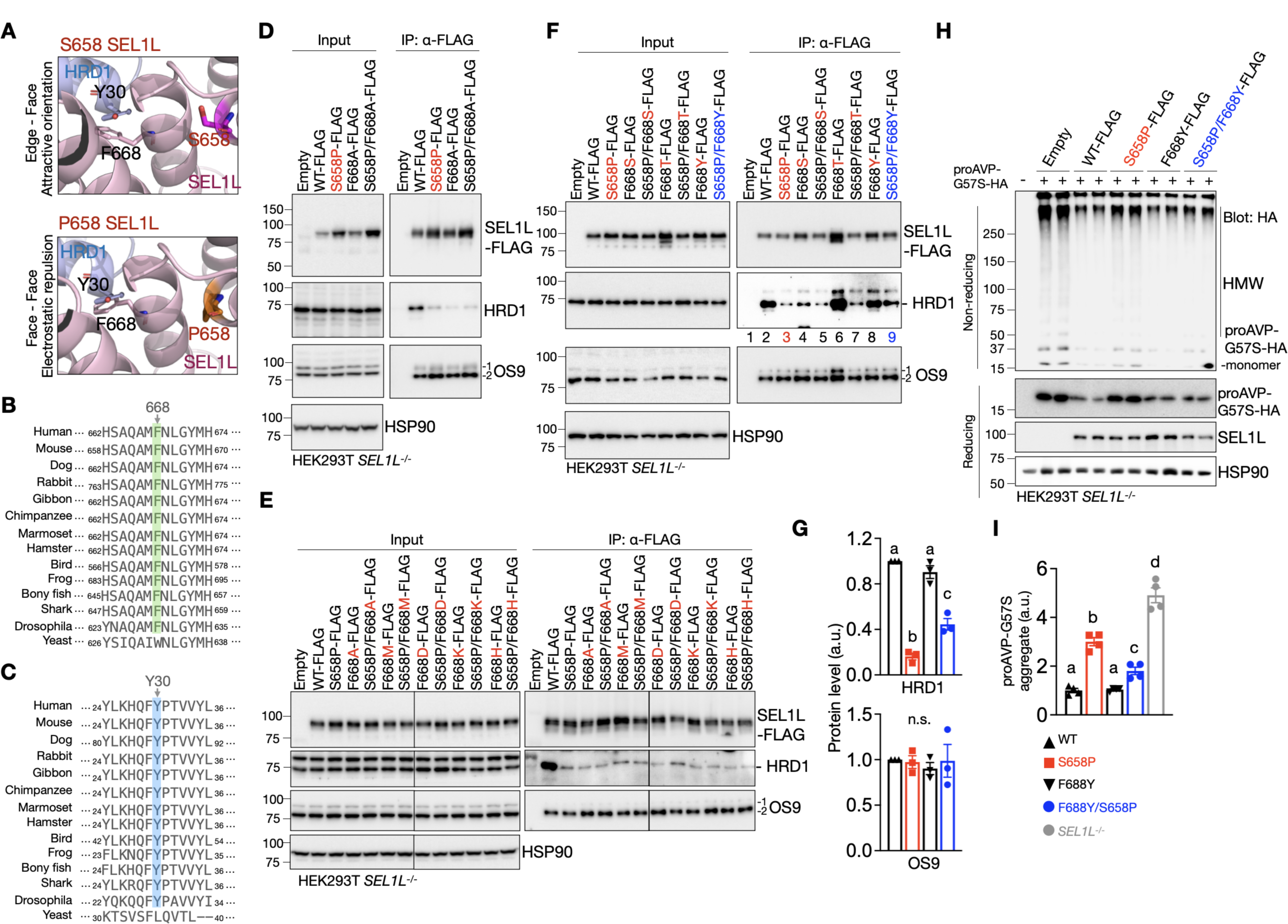
SEL1L S658P disrupts its interaction with HRD1 via SEL1L F668-HRD1 Y30. **(A)** Side views of the aromatic-aromatic interaction of SEL1L F668-HRD1 Y30 in SEL1L-WT (upper) and S658P. **(B-C)** Amino acid sequence alignments of SEL1L and HRD1 showing the conservation of SEL1L-F668 (B, highlighted in green) and HRD1-Y30 (C, highlighted in blue) residues across species. **(D-F)** Immunoprecipitation of FLAG-agarose in *SEL1L^-/-^* HEK293T cells transfected with indicated SEL1L-FLAG constructs to examine the interaction with HRD1 and OS9 (n = 2-3 independent samples for each genotype). **(G)** Quantitation of (F) (n = 3 independent samples for each genotype). **(H-I)** Reducing and non-reducing SDS-PAGE and Western blot analyses of proAVP-G57S high molecular-weight (HMW) aggregates in WT or *SEL1L^-/-^* HEK293T cells transfected with indicated SEL1L-FLAG constructs, with quantitation of proAVP-G57S-HA HMW shown in (I) (n = 4 independent samples for each genotype). Value, mean ± SEM. n.s., not significant; Different letters indicate significant differences at the p=0.05 level by one-way ANOVA followed by Tukey’s post hoc test (G, I).

To test this model experimentally, we asked whether SEL1L^S658P^ disrupts the SEL1L-HRD1 interaction via F668-Y30 interaction. Both SEL1L-F668 and HRD1-Y30 residues are highly conserved from Drosophila to humans, but not in yeast (Figures 6B and 6C). We then mutated F668 to Ala (F668A), which disrupted the SEL1L-HRD1 interaction similarly to S658P and did not rescue the S658P SEL1L-HRD1 interaction (Figures 6D and S6B). Similarly, mutations to other residues including Met (M), Asp (D), Lys (K), His (H), Ser (S) Thr (T), Cys (C) also disrupted the SEL1L-HRD1 interaction and had no effect on rescuing the S658P SEL1L-HRD1 interaction (Figures 6E, 6F and S6C).

Moreover, while mutations to Tyr (Y), Trp (W) or Asn (N) did not affect the interaction between SEL1L and HRD1, only F668Y partially rescued the S658P SEL1L-HRD1 interaction by 2 folds (lane 9 vs. 3, Figures 6F, 6G and S6C). None of them affected the interaction between SEL1L and OS9 (Figures 6D-6G, S6B and S6C). We then asked whther the resuced interacion in SEL1L-F668Y/S658P may improve HRD1 ERAD function. Indeed, unlike SEL1L S658P, SEL1L S658P/F668Y partially reversed the accumulation and HMW aggregation of proAVP-G57S protein upon transfected into *SEL1L^-/-^* HEK293T cells, albeit to a lesser extent than those of SEL1L WT and F668Y (Figures 6H and 6I). These data suggest that SEL1L S658P causes a physical collision between SEL1L F668 and HRD1 Y30, thereby attenuating SEL1L-HRD1 interaction.

### SEL1L-HRD1 interaction is required in forming a functional HRD1 ERAD complex

To gain further insights into the role and importance of SEL1L in HRD1 ERAD, we performed unbiased proteomics LC-MS analyses to map SEL1L and HRD1 interactomes in WT and *SEL1L^-/-^*, WT and *HRD1^-/-^* HEK293T cells, respectively. Following validation of HRD1 and SEL1L antibodies for IP (Figures S7A and S7B), we performed three independent repeats of SEL1L- and HRD1-IP-MS, applied stringent criteria (Figures S7C and S7D) and identified 24 and 47 high-confidence hits for SEL1L and HRD1 interacting proteins, respectively (Figures 7A and 7B, Tables S1 and S2). In line with previous studies ^22, 23^, most of the known components of the SEL1L-HRD1 ERAD complex, including HRD1, SEL1L, OS9, ERLEC1, UBE2J1, DERL2, HERP1, FAM8A1 and VCP (Figure S7E), were identified (Tables S1 and S2), validating our experimental system. We next further interrogated the HRD1-interactome (24 hits) into two groups based on whether the interaction was SEL1L dependent (Group I) or not (Group II) (Figures 7C and 7D).

**Figure 7.**
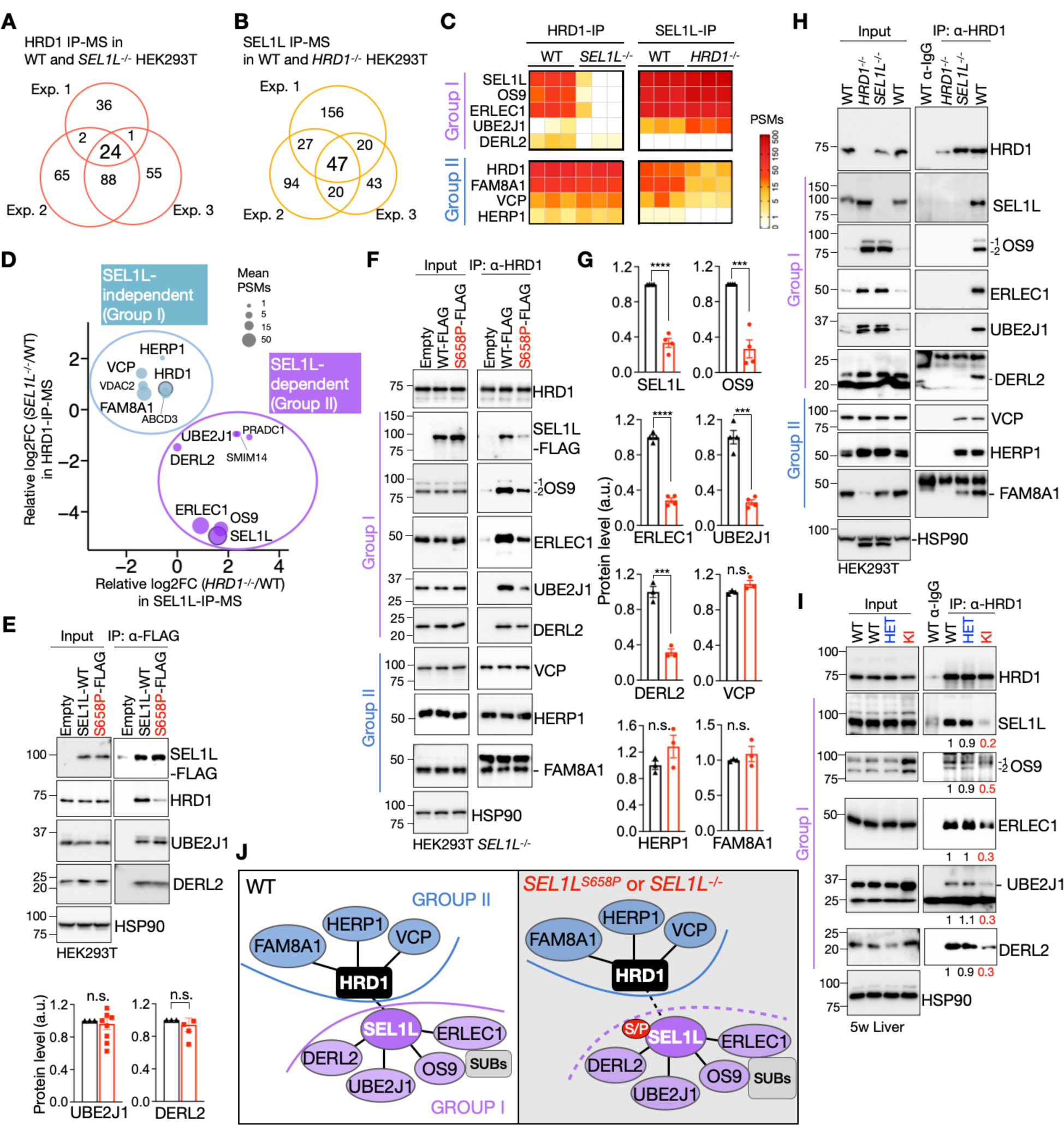
SEL1L-HRD1 interaction is a prerequisite for a functional HRD1 ERAD complex. **(A-B)** Venn diagram showing HRD1- and SEL1L-interacting proteins identified in three independent experiments of HRD1-(A) and SEL1L-(B) IP-MS. **(C)** Heatmaps of known SEL1L-HRD1 ERAD components from the IP-MS experiments. The proteins were grouped into I and II based on their dependency to SEL1L. **(D)** Scatter plot showing average log2 fold change (FC) of the PSMs in *HRD1^-/-^* cells compared to WT in SEL1L-IP-MS and of the PSMs in *SEL1L^-/-^* cells compared to WT in HRD1-IP-MS for the 16 overlapping proteins, plus DERL2. Size of the dots is proportional to the mean PSMs in WT samples. **(E)** Immunoprecipitation of FLAG-agarose in *SEL1L^-/-^*HEK293T cells transfected with indicated SEL1L-FLAG constructs to examine the interaction with HRD1, UBE2J1 and DERL2, with quantitation of UBE2J1 and DERL2 shown below (n = 4-8 independent samples per genotype). **(F-G)** Immunoprecipitation of HRD1 in *SEL1L^-/-^* HEK293T cells transfected with indicated SEL1L-FLAG constructs to examine the endogenous HRD1 interaction with the ERAD components, with quantitation shown in (G) (n = 3-4 independent samples per genotype). **(H)** Immunoprecipitation of endogenous HRD1 in WT, *HRD1^-/-^*and *SEL1L^-/-^* HEK293T cells to examine its interaction with the ERAD components (two independent repeats). **(I)** Immunoprecipitation of HRD1 in the livers of 5-week-old WT and KI mice to examine the endogenous HRD1 interaction with other ERAD components, with quantitation shown below the gel as means from two independent repeats. **(J)** A model showing the formation of SEL1L-HRD1 complex in WT, *SEL1L*^*S*658*P*^ or *SEL1L^-/-^* cells. S/P indicates S658P. Values, mean ± SEM. n.s., not significant; ***p<0.001 and ****p<0.0001 by two-tailed Student’s *t*-test (E, G).

Lectins OS9 and ERLEC1 expectedly formed a tight cluster with SEL1L, i.e. in Group I, through which they interacted with HRD1 (Figures 7C and 7D). Unexpectedly, UBE2J1 and DERL2 also appeared in Group I, suggesting that their interaction with HRD1 may also be mediated through SEL1L. In yeast, Hrd1p and Der1p proteins form a retrotranslocon for misfolded substrates ^19^. Mammals have three DERLIN proteins, with DERL2 primarily associated with the SEL1L-HRD1 complex ^21, 22, 60^. Consistent with this, DERL2 was the only DERLIN protein identified in our IP-MS. On the other hand, three known ERAD components (HERP1, VCP and FAM8A1) and two new hits (VDAC2 and ABCD3) appeared in the Group II, where their interactions with HRD1 were SEL1L independent (Figures 7C and blue dots, 7D). Nonetheless, knockout VDAC2 or ABCD3 failed to affect HRD1 ERAD function towards known substrates (Figures S7F and S7G). Hence, their role in HRD1 ERAD remained uncertain.

We next experimentally tested whether SEL1L interaction with HRD1 is indeed required to bring in UBE2J1 and DERL2, two known critical components of HRD1 ERAD. To this end, we tested the impact of both *SEL1L*^*S*658*P*^ and *SEL1L^-/-^* in the formation HRD1 functional complex with UBE2J1 and DERL2. While *SEL1L*^*S*658*P*^ attenuated its interaction with HRD1, its interaction with the Group I proteins, namely OS9, ERLEC1 (Figures 5D and 5E), UBE2J1 and DERL2 (Figures 7E), was not affected. On the other hand, HRD1 interactions with these Group I proteins were attenuated by 3-4 folds in *SEL1L*^*S*658*P*^-expressing cells (Figures 7F and 7G) and completely abolished in *SEL1L^-/-^* HEK293T cells (Figure 7H). However, HRD1 interactions with the Group II proteins (e.g. HERP1, VCP and FAM8A1) were unaffected by the SEL1L status (Figures 7F and 7H). In line with the overexpression experiments, HRD1 interactions with Group I, not Group II, proteins were reduced in *SEL1L*^*S*658*P*^ KI livers, compared to those in WT and HET livers (Figure 7I). Hence, we conclude that SEL1L is indispensable for HRD1 ERAD function by recruiting Group I factors to HRD1 to form a functional HRD1 ERAD complex (Figure 7J).

## DISCUSSION

SEL1L-HRD1 ERAD plays a critical role in clearing misfolded proteins in the ER. Individually, ample studies have shown that they are indispensable *in vivo*. However, one key question in the field is whether they always work together or have their own function other than HRD1 ERAD, or in the case of HRD1, whether it can function without SEL1L. Here we provide definitive evidence, powered by *in vivo* mouse models, rigorous mutagenesis and proteomic analyses, that SEL1L interaction with HRD1 is pathologically important as a partial loss of the interaction causes disease in mice and dogs, and is biochemically significant as this interaction is critical to form a complete functional HRD1 ERAD complex by bringing in not only lectins, but also the E2 UBE2J1 and retrotranslocon DERLIN. In other words, SEL1L is not only important for HRD1 protein stability ^5, 9, 10^, but also required for substrate recruitment, ubiquitination and retrotranslocation in HRD1 ERAD, essentially all aspects of HRD1 ERAD (Figure 7J). In the absence of SEL1L-HRD1 interaction, HRD1 becomes insufficient with impaired ERAD function (Figure 7J). These unexpected findings not only demonstrate the pathological significance of this ERAD complex in a whole organism (rather than cell type-specific mouse models), but also offer novel insights into the assembly of a functional SEL1L-HRD1 ERAD complex. This study will provide new framework for future therapeutic designs aiming at targeting the SEL1L-HRD1 interaction and function using small molecules.

The importance of SEL1L-HRD1 interaction *in vivo* remained unknown. Here we show that defects merely in SEL1L-HRD1 interaction are sufficient to drive disease pathogenesis including partial embryonic lethality, developmental delay, microcephaly and early-onset ataxia in mice as well as in Finnish Hounds as previously reported ^54^. The *SEL1L*^*S*658*P*^ KI mice suffering from early-onset ataxia are associated with a mild reduction in Purkinje cell numbers. Purkinje cells are some of the largest neurons in the brain with a prominent ER structure that form an interconnected network regulating calcium signaling, protein synthesis, folding, and protein trafficking ^61^. Previous studies have shown that Purkinje cells are vulnerable to alterations in ER homeostasis, as they have a high demand for protein synthesis to maintain a large number of synapsis ^62, 63^. However, as the *SEL1L*^*S*658*P*^ variant is only associated with very mild ER stress in the brain, we speculate that its impact on Purkinje cells as well as ataxia pathogenesis may arise through substrate(s)-related mechanisms, rather than ER stress. The nature of the substrates in these neurons requires future careful investigation. Providing further support for the subtle UPR associated with the variant, examination of peripheral tissues of KI mice including the pancreas, livers, kidneys and adipose tissues revealed no obvious histological or functional defects. The lack of UPR in the KI mice is likely due to various adaptive mechanisms in response to a hypomorphic variant, including, but not limited to, the upregulation of ER chaperones to increase folding efficiency, enhanced aggregation and sequestration of misfolded proteins to hence attenuate proteotoxicity and the activation of ER-phagy to clear misfolded protein aggregates in the ER ^64, 65^.

Previous studies of mammalian SEL1L-HRD1 ERAD complex using overexpression systems have showed that overexpressing SEL1L or HRD1 can pull down OS9, ERLEC1, UBE2J1, HERP1, FAM8A1 and DERL2 ^20–23^. One study showed that SEL1L knockdown did not impact the HRD1-UBE2J1 interaction, while loss of HRD1 disrupts the SEL1L-UBE2J1 connection, hence concluding that HRD1 directly interacts with UBE2J1 ^20^. While our study corroborates some of these findings, our results further identified proteome-wide SEL1L-dependent HRD1 interactors which surprisingly include UBE2J1 and DERL proteins. Indeed, HRD1 interaction with UBE2J1 and DERL was attenuated in cells expressing *SEL1L*^*S*658*P*^ variant or entirely abolished in *SEL1L^-/-^* cells, hence dramatically expanding the role and significance of SEL1L in HRD1 ERAD.

Using a combination of AI structural prediction and biochemical analyses, we showed that SEL1L^S658P^ attenuates the SEL1L-HRD1 interaction via an aromatic-aromatic interaction of SEL1L F668 and HRD1 Y30. While the structure of the SEL1L-HRD1 complex is similar to the yeast Hrd3p-Hrd1p complex (Figure S5C), none of these residues is present in yeast Hrd3p. This finding suggests that, contrary to the yeast Hrd3p-Hrd1p interaction, mammalian SEL1L may play an additional role in the SEL1L-HRD1 interaction. Intriguingly, our data showed that F668W mutation partially rescues the S658P SEL1L-HRD1 interaction, potentially providing a therapeutic target for the treatment of ERAD-associated diseases associated with attenuated SEL1L-HRD1 interaction in the future.

## AUTHOR CONTRIBUTIONS

L.L.L. designed, performed most experiments and wrote the methods and figure legends; X.W. performed the structural analysis; Y.L. performed the proteomics analysis; H.H.W., B.P., M.T., Z.J.L., H.M. and H.W. assisted with some *in vitro* and *in vivo* experiments; X.L. and Z.Z. performed MRI, involved in experimental design and provided insightful discussion; L.Q. and S.S. directed the study and wrote the manuscript with L.L.L; all authors commented on and approved the manuscript.

## ACKNOWLEDGEMENTS

We thank Dr. Chih-Chi Andrew Hu (Houston Methodist Hospital) for reagents; Drs. Ingrid Bergin and Kendra Andrie at the ULAM In-Vivo Animal Core at University of Michigan Medical School for tissue pathological analyses; the Molecular Genetics Core of the Michigan Diabetes Research Center and the Transgenic Animal Model Core for transgenic animal production; the Proteomics Resource Facility, the Microscopy and Image Analysis Core (NIH S10OD28612-01-A1 and P30DK20572), animal phenotyping cores (NIH P30AR069620, P30DK020572, P30DK089503 and 1U2CDK110678-01), and the ULAM In-Vivo Animal Core at University of Michigan Medical School for assistance; and members of the Qi and Arvan laboratories for technical assistance and insightful discussions. This work was supported by RF1NS122060 (Z.Z.), R01DK128077, R01DK132068 (S.S.), R01DK120047, R01DK120330, R35GM130292 and Michigan Protein Folding Disease Initiative (L.Q.). L.L.L. and Z.J.L. are supported in part by National Ataxia Foundation Post- and Pre-doctoral Fellowships (NAF 918037 and 1036307).

## Data and materials availability

The materials and reagents used are either commercially available or available upon the request. Proteomics data will be deposited into a public database PRIDE. All other data are available in the main text or in the Supplemental Information.

