## Supplemental Figures for "SEL1L-HRD1 interaction is prerequisite for the formation of a functional HRD1 ERAD complex"

**This PDF file includes:**

- Key resources table
- Methods
- Supplemental Figures 1-7
- Video S1
- Table S1 and S2
- References

### MATERIALS AND METHODS

### KEY RESOURCES TABLE

| REAGENT or RESOURCE | SOURCE | IDENTIFIER |
|---|---|---|
| **Antibodies** | | |
| Rabbit polyclonal anti-SEL1L | (Zhou, Z., et al)[1] | Epitope hSEL1L 23 -194 aa |
| Rabbit polyclonal anti-HRD1 | Proteintech | Cat# 13473-1-AP |
| Rabbit polyclonal anti-HRD1 | Cell Signaling | Cat# 14773 |
| Rabbit polyclonal anti-HRD1 | This paper | Epitope hHRD1 310 - 550 aa |
| Rabbit polyclonal anti-OS9 | Abcam | Cat# ab109510 |
| Rabbit monoclonal anti-ERLEC1 | Abcam | Cat# ab181166 |
| Rabbit polyclonal anti-DERL2 | Dr. Andrew Hu, Houston Methodist Research Institute | NA |
| Mouse monoclonal anti-UBE2J1 | Santa Cruz | Cat# sc-377002 |
| Rabbit polyclonal anti-VCP | Proteintech | Cat# 10736-1-AP |
| Rabbit polyclonal anti-HERP1 | Abcam | Cat# ab150424 |
| Rabbit polyclonal anti-FMA8A1 | Proteintech | Cat# 24746-1-AP |
| Rabbit polyclonal anti-VDAC2 | Proteintech | Cat# 11663-1-AP |
| Mouse monoclonal anti-ABCD3 | Proteintech | Cat# 66697-1-Ig |
| Rabbit polyclonal anti-CD147 | Proteintech | Cat# 11989-1-AP |
| Rabbit polyclonal anti-IRE1α | Cell Signaling | Cat# 3294 |
| Rabbit polyclonal anti-Caspase-3 | Cell Signaling | Cat# 9665 |
| Rabbit polyclonal anti-Cleaved-caspase-3 | Cell Signaling | Cat# 9661 |
| Rabbit polyclonal anti-PERK | Cell Signaling | Cat# 3192 |
| Rabbit monoclonal anti-p-Thr980 PERK | Cell Signaling | Cat# 3179 |
| Rabbit polyclonal anti-eIF2α | Cell Signaling | Cat# 9722 |
| Rabbit polyclonal anti-p-Ser51 eIF2α | Cell Signaling | Cat# 9721 |
| Rabbit polyclonal anti-BiP/GRP94 | Abcam | Cat# ab21685 |
| Rabbit polyclonal anti-PDI | Enzo | Cat# ADI-SPA-890 |

| | | |
|---|---|---|
| Mouse monoclonal anti-HSP90 | Santa Cruz | Cat# sc-13119 |
| Mouse monoclonal anti-FLAG | Sigma | Cat# F1804 |
| Mouse monoclonal anti-HA | Sigma | Cat# H3663 |
| Rabbit polyclonal anti-Myc | Sigma | Cat# C3956 |
| Rabbit monoclonal anti-Calbindin | Cell Signaling | Cat# 2173 |
| Mouse monoclonal anti-KDEL | Novus Biologicals | Cat# 97469 |
| guinea pig polyclonal anti-NeuN | Sigma | Cat# ABN90 |
| Mouse monoclonal anti-GFAP | Cell Signaling | Cat# 3670 |
| Rabbit polyclonal anti-Iba1 | Fujifilm | Cat# 019-19741 |
| Goat Anti-Rabbit IgG (H + L)-HRP | Bio-rad | Cat# 1706515 |
| Goat Anti-Mouse IgG (H + L)-HRP | Bio-rad | Cat# 1706516 |
| Anti-rabbbit TrueBlot-HRP | Rockland | Cat# 18-8816-33 |
| Anti-mouse TrueBlot-HRP | Rockland | Cat# 18-8817-31 |
| Normal Rabbit IgG | Cell Signaling | Cat# 2729 |
| **Chemicals, Peptides, and Recombinant Proteins** | | |
| DMEM | Fisher | Cat# 092294 |
| Triton X-100 | Fisher | Cat# BP151 |
| Nonidet P-40 | VWR | Cat# M158 |
| Methanol | Fisher | Cat# A433F-1GAL |
| Acetic acid | ThermoFisher | Cat# AC423220025 |
| Paraformaldehyde | Sigma | Cat# 158127 |
| HyClone Fetal Bovine Serum | Fisher | Cat# SH30910.03 |
| Sorenson’s buffer | EMS | Cat# 11600-05 |
| Paraformaldehyde | EMS | Cat# 15710 |
| Glutaraldehyde | EMS | Cat# 16120 |
| TRI Reagent | Molecular Research Center | Cat# TR 118 |
| BCP | Molecular Research Center | Cat# BP 151 |
| Protease Inhibitor Cocktail | Sigma | Cat# P8340 |
| Phosphatase inhibitor cocktail II | Sigma | Cat# P5726 |

| | | |
|---|---|---|
| Pierce BCA protein assay kit | ThermoFisher | Cat# 23225 |
| Lipofectamine 2000 | Invitrogen | Cat# 11668019 |
| Polyethylenimine | Fisher | Cat# 2397661 |
| Opti-MEM | ThermoFisher | Cat# 31985070 |
| Cycloheximide | Calbiochem | Cat# 239764 |
| PfuUltra DNA Polymerase | Agilent Technologies | Cat# 600380 |
| SuperScript III Reverse Transcriptase | Invitrogen | Cat# 18080093 |
| DAPI | Vector Laboratories | Cat# H-1500 |
| Anti-FLAG M2 affinity gel | Sigma | Cat# A2220 |
| Anti-Myc agarose | Sigma | Cat# 7470 |
| Protein A agarose | Invitrogen | Cat# 20333 |
| **Experimental models: cell lines** | | |
| HEK293T | ATCC | CRL-3216 |
| **Experimental models: organisms/strains** | | |
| Mouse: Sel1L S658P knock-in (KI); B6.SJL | This paper | NA |
| **Recombinant DNA** | | |
| mSEL1L(WT)-FLAG | This paper | NA |
| mSEL1L(mutation)-FLAG | This paper | NA |
| hHRD1-Myc | Gifted from Dr. Yihong Ye | NA |
| proAVP(G57S)-HA | (Guo, S., et al)[2] | NA |
| lentiCRISPR-hHRD1 | This paper | NA |
| lentiCRISPR-hSEL1L | This paper | NA |
| lentiCRISPR-hABCD3 | This paper | NA |
| lentiCRISPR-hVDAC2 | This paper | NA |
| **Software and Algorithms** | | |
| Fiji | Fiji | Fiji.sc |
| GraphPad Prism 8 | GraphPad | Graphpad.com |

| | | |
|---|---|---|
| Image Lab | Bio-rad | Bio-rad.com |
| PyMOL | PyMOL | pymol.org |

### Mice

The $SEL1L^{S658P}$ knock-in (KI) mice (human SEL1L p.Ser658 is equivalent to mouse SEL1L p.Ser654) were generated at the University of Michigan Molecular Genetics Core using the CRISPR-Cas9 technology. The two single guide RNAs (sgRNAs) were designed by using computer algorithm (http://crispor.tefor.net), targeting mouse genome *SEL1L* exon 19 where the mutation is located: sgRNA1: 5'-CAGGCGGTAATGAATAAATGCGG-3'; sgRNA2: 5'-CTGGGCTACATGCACGAGAAGGG -3'. The sgRNAs were synthesized using the Synthego sgRNA synthesis Kit per manufacturer's protocol, tested and confirmed in fertilized eggs. A donor DNA carrying mouse *Sel1L* cDNA 1960C>T mutation and additional silent mutations was designed to mediate homology-directed repair (HDR): 5'-GTTACACTGTGGCTAGAATTAAGCTTGGAGACTACCACTTCTATGGCTTTGGCACTGATGTGGATTATGAGACAGCCTTCATTCACTACCGGCTGGCTCCTGAGCAGCAGCACAGCGCCCAAGCTATGTTTAACCTGGGATACATGCATGAGAAAGGCCTAGGCATTAAACAGGTGAGTGTGGGCCACGCTGGCGCTGAG-3'. The donor DNA was synthesized as Ultramer dsDNA by Integrated DNA Technologies, Inc. A mixture of Cas9 protein (Sigma), sgRNAs, and donor DNA was microinjected into fertilized mouse eggs on the B6/SJL background. The injected zygotes were then transferred into pseudopregnant females. A pair of primers were designed for genotyping: Ser658Pro-F (5'-ATTCACTACCGGCTGGCTC-3') and WT-R (5'-TCCTGTAGCCCGAGGTCTGA-3'). Tail samples were obtained from 2-week-old pups for genotyping and heterozygous $SEL1L^{S658P}$ KI mice were established as F0 founders. The existence of desired mutations was further confirmed by Sanger sequencing in two independent founders. The founders were then bred separately to WT C57BL6/J mice to obtain F1 heterozygous $SEL1L^{S658P}$ KI mice. F1 heterozygous $SEL1L^{S658P}$ mice were inter-crossed to generate homozygous $SEL1L^{S658P}$ KI mice and its WT and heterozygous littermates.

To study embryonic lethality, pregnant heterozygous female mice were euthanized at embryonic day 10-12 (E10-12) or at E14-16. Uterine horns were removed and placed them in a petri dish filled with cold PBS. The embryos were removed from the amniotic membrane and transferred into a new petri filled with cold PBS. A small tail biopsy was used for genotyping of individual embryos. The post-natal lethality data were obtained from 71 litters of 28 breeding pairs total.

Age- and gender-matched littermates were maintained in a temperature-controlled room on a 12-h light/dark cycle and used in all studies. All animal procedures were approved by the Institutional Animal Care and Use Committee of the University of Michigan Medical School (PRO0008989 and PRO00010658) and in accordance with the National Institutes of Health (NIH) guidelines.

### Genotyping

Mice were routinely genotyped using PCR of genomic DNA samples obtained from tails or ears with the following primer pairs:

*SEL1L*$^{S658P}$ allele for *SEL1L*$^{S658P}$ KI mice:

F: 5'-ATTCACTACCGGCTGGCTC-3';

R: 5'-TCCTGTAGCCCGAGGTCTGA-3';

*SEL1L* wildtype allele for *SEL1L*$^{S658P}$ KI mice:

F: 5'-CGCATTTATTCATTACCGCCTG-3';

R: 5'-TCCTGTAGCCCGAGG TCTGA-3';

### Behavioral studies

All behavior procedures were performed by investigators blind to the genotype of each group or nature of intervention. The indirect calorimetry (including food intake) was performed and analyzed at University of Michigan Animal Phenotyping Core as previously described [3,4]. Food intake was measured in single-housed mice on a regular chow diet at room temperature (20-23 °C). The measurements were carried out continuously for 72 hours.

For the hindlimb clasping assessment, mice were lifted up by tails and held over a cage for 1 min to assess abnormal hind limb clasping and scored as previously described [5]. For the hindlimb clasping assessment, mice were lifted by their tails and held over a cage for 1 min. The mice were then given a score ranging from 0 to 4 based on the following criteria:

0 - No limb clasping

1 - One hindlimb splayed incompletely; toes splayed normally

2 - Both hindlimbs splayed incompletely; toes splayed normally

3 - Both hindlimbs clasped and both forelimbs clasped; toes curled

4 - Forelimbs and hindlimbs all clasped together; toes curled

For the gait analysis, hind- and fore-paws were coated with red and black nontoxic food coloring, respectively. Mice were then allowed to walk along a 450-mm-long, 50-mm-wide runway with 70-mm-high walls with white paper lining the floor, into a darkened enclosed escape box. The footprints were analyzed for stride length (the distance covered by the same hind paw), the stride width (the distance from one hind-limb that intersects perpendicularly with the line for stride length on the contralateral hind paw), paw matched distance (the distance between hind and forepaw) and the ratio between stride length and width. All mice received both three trainings and trial run.

The balance beam study was used to evaluate motor coordination and balance as previously described [6]. Mice were trained to stay upright and walk across an elevated narrow beam to a safe platform for two consecutive days (three times per day). The beam apparatus consisted of a flat beam that was 1-meter long and 12 mm wide, resting 50 cm above the table on top of two poles. A darkened escape box with nesting material was placed at the end of the beam as the finish point. On the third day, the time to cross a distance of 80 cm located in the center of the beam was measured. A video camera was set on a tripod to record the performance of the animals during the test. Room temperature, humidity, lighting, and background noise were kept consistent throughout the experiment.

For rotarod test, mice were placed into a rod rotating (Model LE8500, Panlab SL) at an accelerating speed. Animals were tested in four trials per day for three consecutive days. The tests started with 4 rpm and increased to 40 rpm after 300 sec, the time and the maximal speed until mice fell from the rod were measured.

#### Histology and immunofluorescence

Anesthetized mice were perfused with 20 ml of 0.9% NaCl followed by 40 ml of 4% paraformaldehyde in 0.1 M PBS pH 7.4 for fixation. Tissues were dissected out and fixed overnight in 4% paraformaldehyde in PBS at 4ºC. For hematoxylin and eosin (H&E) staining, samples were dehydrated, embedded in paraffin, and stained at the Rogel Cancer Center Tissue and Molecular Pathology Core or In-Vivo Animal Core at the University of Michigan Medical School. Quantitation of Purkinje cell numbers was performed on H&E stained sections. Counts were normalized to the length of the Purkinje cell layer, as measured by Aperio ImageScope software, and reported as Purkinje cell density. For immunofluorescence staining, paraffin-embedded brains sections were deparaffinized in xylene and rehydrated using graded ethanol series (100%, 90%, 70%), followed by rinse in distilled water. Antigen retrieval was performed by boiling the slides in a microwave in sodium citrate buffer. Sections were then incubated in a blocking solution (5% donkey serum, 0.3% Triton X-100 in PBS) for 1 hour at room temperature and with primary antibodies (Calbindin, Cell Signaling, #2173, 1:100; KDEL, Novus Biologicals, #97469, 1:200; NeuN, Sigma, #ABN90, 1:200; GFAP, Cell Signaling, #3670, 1:100; Iba1, Fujifilm, #019-19741, 1:100) overnight at 4°C in a humidifying chamber. The next day, following 3 washes with PBST (0.03% Triton X-100 in PBS), slides were incubated with the respective Alexa Fluor–conjugated to secondary antibodies (Jackson ImmunoResearch; dilution 1:500) for 1 hour at room temperature, followed by mounting with VECTASHIELD mounting medium containing DAPI (Vector Laboratories, H-1500). Images were captured using a Nikon A1 confocal microscope at the University of Michigan Morphology and Image Analysis Core.

#### Serum chemistry tests

Mouse blood was collected without anticoagulation via cardiac puncture immediately prior to sacrifice in mice anesthetized by isoflurane inhalation. The whole blood was allowed to clot under room temperature for 15-30 min and then serum was separated by centrifugation. Serum Mini Chemistry Panel Tests (Alanine Transaminase, Total Bilirubin, Alkaline Phosphatase, Blood Urea Nitrogen, Creatinine) were performed at the Unit for Laboratory Animal Medicine (ULAM) In Vivo Animal Core pathology laboratory at the University of Michigan.

#### 3D MRI brain analysis

Anesthetized mice were transcardially perfused with 4% paraformaldehyde in PBS. The head were cut and removed the skin and soft tissue and then soaked in PBS containing 5% Gd-DTPA and stored in 4°C for 5 days. Prior to imaging, the brain specimens were placed in a 15 ml tube containing proton signal-free susceptibility-matched fluid (Galden Heat Transfer Fluid, HT230. SOLVEY, Italy), which was placed in a mouse head coil. MRI acquisitions were performed on a 7T simultaneous PET-MR scanner (MR Solutions Ltd.) at the Zilkha Neurogenetic Institute (University of Southern California). The 2D sagittal T2-weighted images (T2WI) were performed using fast spin echo (FSE) sequence (TR/TE: 4000/45 ms, echo train length of 7, eight average, 28 slices, slice thickness of 500 μm, in-plane resolution 390×140 $\mu m^3$), to manually draw the regions of interests (ROIs) of cerebellum and cortex. 3D T2Wis were performed using Fast Low Angle Shot (FLASH) sequence (TR/TE: 50/5 ms, FA=30°, seven average, with an isotropic voxel resolution of 93×93×93 $\mu m^3$) for 3D brain render reconstruction For the reconstruction, 3D images were performed using our in-house MATLAB code. The intensity inhomogeneity of images was corrected by N4 algorithm and then co-registered to the Mouse Magnetic Resonance Microscopy Atlas (https://www.loni.usc.edu/research/atlas_downloads). The masks of whole brain, cerebellum and cortex were segmented and manually edited using the ITK-SNAP software. The rendered 3D brain was reconstructed, and masks of cerebellum and cortex were displayed using the ParaView software version 5.10.1.

#### CRISPR/Cas9-based knockout (KO) and KI HEK293T cells

HEK293T cells were cultured at 37°C with 5% $CO_2$ in DMEM with 10% fetal bovine serum (Fisher Scientific). To generate SEL1L-, HRD1-, ABCD3-, VDAC2-deficient HEK293T cells, sgRNA oligonucleotides designed for human *SEL1L* (5'-GGCTGAACAGGGCTATGAAG-3'), human *HRD1* (5'-GGACAAAGGCCTGGATGTAC-3'), human *ABCD3* (5'-GATCAAGTGATATATCCAGA-3', or 5'-GTTATTGTCATCTTACCAAT-3') or human VDAC2 (5'-GAGAGTGTAAGCTTCACACC-3', 5'-GTGCCAAATCAAAGCTGACA-3' or 5'-GATGGGAAGAGCATTAATGC-3') was inserted into lentiCRISPR v2 (plasmid 52961; Addgene). Cells grown in 10 cm petri dishes were transfected with indicated plasmids using 5 µl 1 mg/ml polyethylenimine (PEI, Sigma) per 1µg of plasmids for HEK293T cells. Cells were cultured 24 hours after transfection in medium containing 2 µg/ml puromycin for 24 hours and then in normal growth media.

*SEL1L*$^{S658P}$ KI HEK293T cells was generated using the CRISPR-Cas9 system and Homology-Directed Repair (HDR) mechanism (Integrated DNA Technologies, IDT). 5 µl 100 µM Alt-R crRNA (IDT) with gRNA sequence was first mixed with 5 µl 100 µM Alt-R tracrRNA (IDT) containing Cas9 interacting sequence. To anneal the oligos, the duplex mixture was heated at 95°C for 5 min and then cooled at room temperature for 20 min. 9 µl of the guide complex was incubated with 6 µl 62 µM Alt-R Cas9 enzyme (IDT) at room temperature for 20 min. 5 µl of the ribonucleoprotein (RNP) complex, together with 1.2 µl 100 µM HDR Donor Oligo (IDT), 1.2 µl 100 µM Alt-R Cas9 Electroporation Enhancer (IDT), was mixed with 100 µl HEK293T cell suspension (about $5x10^5$ cells) in Electroporation Solution (Ingenio). The mixture was transferred into a 0.2 cm cuvette and electroporation was performed using Lonza Nucleofector IIb (Lonza). To prepare the cell culture media, 3.4 µl 0.69 mM Alt-R HDR Enhancer V2 (IDT) was added to 2000 µl DMEM with 10% fetal bovine serum (Fisher Scientific). After electroporation, the cell suspension was added to the cell culture media, and the mixture was incubated in 4 wells of a 24-well dish (500 µl per well). The cells were cultured at 37°C with 5% CO2. After 5 days of incubation, the genomic DNA of the cell culture was extracted with 50 mM

NaOH. DNA fragments covering the target sites were amplified by PCR, using HotStart Taq 2X PCR Master (ABclonal), and analyzed by Sanger Sequencing (Eurofins) to estimate the percentage of mutant allele in the cell pool. In parallel, the cell culture was diluted into 8 cells per mL and cultured in eight 96-well plates (100 µl per well) for single cell isolation. After 10 days, 100 single cell colonies were transferred into 24-well plates. The $SEL1L^{S658P}$ region was amplified using a 25 µl PCR reaction and sequenced. Cell colonies with homozygous $SEL1L^{S658P}$ alleles were transferred into a 6-well plate for further experiments.

crRNA (guide sequence): 5'- GTTGCTGCTCAGAAGCCAGA-3'
HDR Donor Oligo (the mutation site is underlined): 5'-
CCCAGATTAAACATAGCTTGTGCACTGTGTTGCTGCTCAGGAGCCAGACGGTAATGAATAA
ATGCAGTTTCATAATCTACATCGG-3'
Amplification PCR primers:
F: 5'-TGTAAGCATGTCAATGGGAGGAG-3'
R: 5'- AGCGTGGATAGCAGTATCTGT-3'
Sequencing primers:
5'- GTTTAAGGCTATACTGTGGCT-3'

#### Protein structure modeling

The individual structure models of human SEL1L, HRD1, OS9, and DERLIN1 were downloaded from AlphaFold2 database (https://alphafold.ebi.ac.uk/) [7]. The complex structure was constructed using TM-align [8] by superposing the predicted human SEL1L, HRD1, OS9, and DERLIN1 structures to the CryoEM structure of the yeast Hrd3p-Hrd1p-Der1p protein complex (PDB ID: 6VJZ) and Hrd3p-Yos9 complex (PDB ID: 6VK3). All the structure images of human SEL1L 171-723 aa, HRD1 1-334 aa, OS9 33-655 aa and DERLIN1 1-213 aa were rendered by PyMOL (version 2.3.2). To analyze the evolutionary conservation of each residue, a position-specific scoring matrix (PSSM) was generated from a PSI-BLAST search of the target protein through the NCBI NR database [9,10].

The amino acid sequence of human SEL1L (accession no. NP_005056.3) was aligned with the SEL1L homologs from chimpanzee (JAA44458.1), gibbon (XP_003260889.1), marmoset (XP_002754215.1), dog (XP_038530327.1), rabbit (XP_002719662.2), hamster (NP_001268291.1), mouse (NP_001034178.1), bird (XP_021143193.1), frog

(XP_041430335.1), bony fish (NP_001038629.1), shark (XP_048393621.1), drosophila (NP_001262882.1) and yeast (QHB10358.1) by using ClustalW program.

The amino acid sequence of human HRD1 (SYVN1) (accession no. NP_115807.1) was aligned with the HRD1 homologs from chimpanzee (JAA39943.1), gibbon (XP_032009276.1), marmoset (JAB46248.1), dog (XP_038280973.1), hamster (XP_051049706.1), mouse (AAH46829.1), bird (XP_053824008.1), frog (XP_012816159.1), bony fish (AAH44465.1), shark (XP_043538543.1), drosophila (NP_001263152.1) and yeast (QHB11597.1) by using ClustalW program.

### Plasmids

The following plasmids were used in the study: (h denotes human genes; m denotes mouse genes): *mSel1L* cDNA was cloned from mouse liver cDNA and inserted into the pcDNA3 to generate pcDNA3-mSEL1L(WT)-FLAG; pcDNA3-h-HRD1-Myc and pcDNA3-h-proAVP(G57S)-HA were described previously [2]; The SEL1L-FLAG mutants S658P, S658A, S658D, S658K, S663P, F668A, F668S, F668T, F668Y, F668M, F668D, F668K, F668H, F668N, F668W and F668C were generated using the plasmid pcDNA3-mSEL1L(WT)-FLAG as the template. P658S and the double mutations of SEL1L (S658P mutated back to WT) was generated using the plasmid pcDNA3-mSEL1L(S658P)-FLAG. All plasmids were validated by DNA sequencing.
The primers are:
mSEL1L-FLAG-F: 5'-CGCGGATCCACCATGCAGGTCCGCGTCAGGCTGTCG-3'
R: 5'-CGCTCTAGACTATTTATCATCATCATCTTTATAATCTCCGCCCTGTG GTGGCTGCTGCTCTGG-3'
S658P-F: 5'-GCATTTATTCATTACCGCCTGGCTCCTGAGCAGC-3'
R: 5'-GCTGCTCAGGAGCCAGGCGGTAATGAATAAATGC
S658A-F: 5'-GCATTTATTCATTACCGCCTGGCTGCTGAGCAGC-3'
R: 5'-GCTGCTCAGCAGCCAGGCGGTAATGAATAAATGC-3'
S658D-F: 5'-GCATTTATTCATTACCGCCTGGCTGATGAGCAGC-3'
R: 5'-GCTGCTCATCAGCCAGGCGGTAATGAATAAATGC-3'
S658K-F: 5'-GCATTTATTCATTACCGCCTGGCTAAAGAGCAGC-3'
R: 5'-GCTGCTCTTTAGCCAGGCGGTAATGAATAAATGC-3'
P658S-F: 5'-GCATTTATTCATTACCGCCTGGCTTCTGAGCAGC-3'
R: 5'-GCTGCTCAGAAGCCAGGCGGTAATGAATAAATGC-3'
S663P-F: 5'-CTTCTGAGCAGCAGCACCCCGCCCAAGCTATG-3'

R: 5'-CATAGCTTGGGCGGGGTGCTGCTGCTCAGAAG-3'
F668A-F: 5'-GCCCAAGCTATGGCTAACCTGGGCTAC-3'
R: 5'-GTAGCCCAGGTTAGCCATAGCTTGGGC-3'
F668S-F: 5'-GCCCAAGCTATGTCTAACCTGGGCTAC-3'
R: 5'-GTAGCCCAGGTTAGACATAGCTTGGGC-3'
F668T-F: 5'-GCCCAAGCTATGACCAACCTGGGCTAC-3'
R: 5'-GTAGCCCAGGTTGGTCATAGCTTGGGC-3'
F668Y-F: 5'-GCCCAAGCTATGTACAACCTGGGCTAC-3'
R: 5'-GTAGCCCAGGTTGTACATAGCTTGGGC-3'
F668M-F: 5'-GCCCAAGCTATGATGAACCTGGGCTAC-3'
R: 5'-GTAGCCCAGGTTCATCATAGCTTGGGC-3'
F668D-F: 5'-GCCCAAGCTATGGATAACCTGGGCTAC-3'
R: 5'-GTAGCCCAGGTTATCCATAGCTTGGGC-3'
F668K-F: 5'-GCCCAAGCTATGAAGAACCTGGGCTAC-3'
R: 5'-GTAGCCCAGGTTCTTCATAGCTTGGGC-3'
F668H-F: 5'-GCCCAAGCTATGCACAACCTGGGCTAC-3'
R: 5'-GTAGCCCAGGTTGTGCATAGCTTGGGC-3'
F668N-F: 5'-GCCCAAGCTATGAATAACCTGGGCTAC-3'
R: 5'-GTAGCCCAGGTTATTCATAGCTTGGGC-3'
F668W-F: 5'-GCCCAAGCTATGTGGAACCTGGGCTAC-3'
R: 5'-GTAGCCCAGGTTCCACATAGCTTGGGC-3'
F668C-F: 5'-GCCCAAGCTATGTGTAACCTGGGCTAC-3'
R: 5'-GTAGCCCAGGTTACACATAGCTTGGGC-3'

#### Western blot and antibodies

Mouse tissues or HEK293T cells were harvested and snap-frozen in liquid nitrogen. The proteins were extracted by sonication in NP-40 lysis buffer (50 mM Tris-HCl at pH7.5, 150 mM NaCl, 1% NP-40, 1 mM EDTA) with protease inhibitor (Sigma), DTT (Sigma, 1 mM) and phosphatase inhibitor cocktail (Sigma). Lysates were incubated on ice for 30 min and centrifuged at 16,000 g for 10 min. Supernatants were collected and analyzed for protein concentration using the Bio-Rad Protein Assay Dye (Bio-Rad). 20-50 μg of protein were denatured at 95°C for 5 min in 5x SDS sample buffer (250 mM Tris-HCl pH 6.8, 10% sodium dodecyl sulfate, 0.05% Bromophenol blue, 50% glycerol, and 1.44 M β-mercaptoethanol). Protein was separated on SDS-PAGE, followed by electrophoretic transfer to PVDF (Fisher

Scientific) membrane. The blots were incubated in 2% BSA/Tri-buffered saline tween-20 (TBST) with primary antibodies overnight at 4°C: anti-HSP90 (Santa Cruz, #sc-13119, 1:5,000), anti-SEL1L (home-made, 1:10,000) [1], anti-HRD1 (Proteintech, #13473-1, 1:2,000), anti-OS9 (Abcam, #ab109510, 1:5,000), anti-CD147 (Proteintech, #11989-1, 1:3,000), anti-IRE1α (Cell Signaling, #3294, 1:2,000), anti-ERLEC1 (Abcam, #ab181166, 1:5,000), anti-UBE2J1 (Santa Cruz, #sc-377002, 1:3,000), anti-DERL2 (gift from Chih-Chi Andrew Hu, 1:1,000), anti-BiP/GRP94 (Abcam, #ab21685, 1:5,000), anti-PDI (Enzo, #ADI-SPA-890, 1:5,000), anti-FLAG (Sigma, #F1804, 1:1,000), anti-HA (Sigma, #H3663, 1:5,000), anti-Myc (Sigma, #C3956, 1:3000), anti-Pro-Caspase-3 (Cell Signaling, #9662, 1:2,000), anti-cleaved-Caspase-3 (Cell Signaling, #9661, 1:1,000), anti-Calbindin (Cell Signaling, #2173, 1:5,000), anti-PERK (Cell Signaling, #3192, 1:5000), anti-p-PERK (Cell Signaling, #3179, 1:1000), anti-eIF2α (Cell Signaling, #9722, 1:5000), anti-p-eIF2α (Cell Signaling, #9721, 1:1000), anti-VCP (Proteintech, #10736-1-AP, 1:3000), anti-HERP1 (Abcam, #ab150424, 1:3000), anti-FAM8A1 (Proteintech, #24746-1-AP,1:3000), anti-VDAC2 (Proteintech, #11663-1-AP,1:3000), anti-ABCD3 (Proteintech, #66697-1-Ig,1:3000). Membranes were washed with TBST and incubated with secondary antibodies, either HRP conjugated (Bio-Rad, 1:10,000), anti-Rabbit IgG TrueBlot HRP (Rockland, #18-8816-33, 1:500) or anti-Mouse IgG TrueBlot-HRP (Rockland, #18-8817-31, 1:500) at room temperature for 1h for ECL chemiluminescence detection system (Bio-Rad) development. Band intensity was determined using Image lab (Bio-Rad) software.

#### Cycloheximide (CHX) treatment of HEK293T cells

To study protein degradation, the HEK293T cells were cultured in the cell culture medium with 50 µg/ml for the indicated time and harvested into liquid nitrogen for protein extraction.

#### Generation of HRD1-specific antibody

Rabbit polyclonal anti-human HRD1 antibody was raised in rabbits immunized with glutathione S-transferase (GST) fused N-terminal 241 amino acids (310 - 550 aa) of human HRD1. Briefly, the cDNA sequence corresponding to the truncated human HRD1 (310 - 550 aa) was sub-cloned into the pGEX-4T-AB1 vector to allow for the expression of the recombinant GST-hHRD1-His protein in *E.coli* Rosetta upon IPTG (0.8 mM) induction at 37°C for 4h. Recombinant GST-hHRD1-His proteins were purified using Ni-NTA agarose. The polyclonal antibody was generated by immunizing rabbits with the recombinant hHRD1 proteins and further affinity-purified using the same antigen.

#### Immunoprecipitation (IP)

HEK293T cells or the livers were snap-frozen in liquid nitrogen and whole cell lysate was prepared in IP lysis buffer [For IP in cells: 150 mM NaCl, 0.2% Nonidet P-40 (NP40), 0.1% Triton X-100, 25 mM Tris-HCl pH 7.5; For IP in liver: 150 mM NaCl, 1% Nonidet P-40 (NP40), 25 mM Tris-HCl pH 7.5] for anti-SEL1L-FLAG, HRD1-Myc, SEL1L and HRD1 IP at 4°C, supplemented with protease inhibitors, protein phosphatase inhibitors, and 10 mM N-ethylmaleimide. A total of ~5 mg protein lysates were incubated with 10 µl anti-FLAG agarose (Sigma, #A2220), 10 µl HA agarose (ThermoFisher, #26182), 10 µl Myc agarose (Sigma, #7470), 1 µl anti-SEL1L home-made antibody or 1 µl anti-HRD1 home-made antibody overnight at 4°C with gentle rocking. SEL1L or HRD1-IP lysates were incubated with Protein A agarose (Invitrogen, #20333) at 4 °C for 2 hours. The incubated agaroses were washed three times with IP lysis buffer and eluted in SDS sample buffer at 95°C for 5 min followed by SDS-PAGE and Immunoblot.

#### (Non-)reducing SDS-PAGE

HEK293T cells transfected with proAVP(G57S)-HA plasmid were snap-frozen in liquid nitrogen and whole cell lysates were prepared in the NP-40 lysis buffer supplemented with protease and phosphatase inhibitors and 10 mM N-ethylmaleimide. Lysates were incubated on ice for 30 min and centrifuged at 16,000 g for 10 min. Supernatants were collected and analyzed for protein concentration using the Bio-Rad Protein Assay Dye (Bio-Rad). For reducing SDS-PAGE analysis, lysates were denatured at 95°C for 5 min in the 5x SDS sample buffer. For non-reducing SDS-PAGE analysis, the lysates were prepared in 5x non-denaturing sample buffer (250 mM Tris-HCl pH 6.8, 1% sodium dodecyl sulfate, 0.05% bromophenol blue, 50% glycerol) and incubated at 37°C for 1 hour. The samples were loaded into a 4%-12% gradient.

#### RNA preparation and RT-PCR

Total RNA was extracted from tissues using TRI Reagent and BCP phase separation reagent (Molecular Research Center, TR 118). The ratio of *Xbp1s* to total *Xbp1* (*Xbp1u* + *Xbp1s*) levels was quantified by Image Lab (Bio-Rad) software. For RT-PCR analysis the following primer sequences were used:

*mXbp1* F: ACGAGGTTCCAGAGGTGGAG R: AAGAGGCAACAGTGTCAGAG
*mL32* F: GAGCAACAAGAAAACCAAGCA R: TGCACACAAGCCATCTACTCA

#### Transmission electron microscopy (TEM)

Mice were anesthetized and perfused with 3% glutaraldehyde, 3% formaldehyde in 0.1 M cacodylate buffer (Electron Microscopy Sciences, 16220, 15710, 11653). Cerebellum was dissected out, cut into small pieces, and fixed overnight at 4ºC in 3% glutaraldehyde, 3% formaldehyde in 0.1 M Sorenson's buffer (Electron Microscopy Sciences, 11682). The tissues were then prepared, embedded, and sectioned at the University of Michigan Histology and Imaging Core. Samples were stained with uranyl acetate/lead citrate and high-resolution images were acquired with a JEOL 1400-plus electron microscope (JEOL).

### Mass spectrometry

For SEL1L- and HRD1-IP in HEK293T cells, immunoprecipitation of endogenous SEL1L or HRD1 in WT, *SEL1L*$^{-/-}$ or *HRD1*$^{-/-}$ HEK293T cells were performed using 10 mg proteins from each sample lysed with the NP-40 lysis buffer supplemented with protease inhibitors, protein phosphatase inhibitors, and 10 mM N-ethylmaleimide. The cell lysates were first incubated anti-SEL1L (home-made) or anti-HRD1 (home-made or Cell Signaling #14773) at 4°C overnight, followed by incubation with Protein A agarose (Invitrogen, #20333) at 4°C for 2 hours. An IP reaction with IgG using cell lysate of *SEL1L*$^{-/-}$ *or HRD1*$^{-/-}$ cells was included as a negative control for SEL1L- or HRD1-IP, respectively.

The beads were resuspended in 50 µl of 0.1 M ammonium bicarbonate buffer (pH~8). Cysteines were reduced by adding 50 µl of 10 mM DTT and incubating at 45° C for 30 min. Samples were cooled to room temperature and alkylation of cysteines was achieved by incubating with 65 mM 2-Chloroacetamide, under darkness, for 30 min at room temperature. An overnight digestion with 1 µg sequencing-grade modified trypsin was carried out at 37° C with constant shaking in a Thermomixer. Digestion was stopped by acidification and peptides were desalted using SepPak C18 cartridges using manufacturer's protocol (Waters). Samples were completely dried using vacufuge. Resulting peptides were dissolved in 0.1% formic acid/2% acetonitrile solution and were resolved on a nano-capillary reverse phase column (Acclaim PepMap C18, 2 micron, 50 cm, ThermoScientific) using a 0.1% formic acid/2% acetonitrile (Buffer A) and 0.1% formic acid/95% acetonitrile (Buffer B) gradient at 300 nl/min over a period of 180 min (2-25% buffer B in 110 min, 25-40% in 20 min, 40-90% in 5 min followed by holding at 90% buffer B for 10 min and re-equilibration with Buffer A for 30 min). Eluent was directly introduced into Q exactive HF mass spectrometer (Thermo Scientific, San Jose CA) using an EasySpray source. MS1 scans were acquired at 60K resolution (AGC target=3x106; max IT=50 ms). Data-dependent collision induced dissociation MS/MS spectra were acquired using Top speed method (3 seconds) following each MS1 scan (NCE ~28%; 15K resolution; AGC target 1x105; max IT 45 ms).

Proteins were identified by searching the MS/MS data against UniProt entries using Proteome Discoverer (v2.4, Thermo Scientific). Search parameters included MS1 mass tolerance of 10 ppm and fragment tolerance of 0.2 Da; two missed cleavages were allowed; carbamidomethylation of cysteine was considered fixed modification and oxidation of methionine, deamidation of asparagine and glutamine were considered as potential modifications. False discovery rate (FDR) was determined using Percolator and proteins/peptides with an FDR of ≤1% were retained for further analysis.

SEL1L- and HRD1-interacting proteins were selected based on the peptide spectrum matches (PSMs) from the label-free IP-MS results. For each protein hit, the PSM value from the IgG sample must be smaller than one-tenth of the WT sample from the same experiment; the PSMs ratio of the bait KO sample to WT sample must be smaller than the ratio of the corresponding bait; specifically for the hits passed selection in two but not all three experiments, a minimal PSM value of 2 in at least one WT sample is reinforced. Common contaminating proteins in IP experiments, including keratin, keratin-associated proteins, ribosomal proteins and nuclear proteins were excluded. The proteins in association with both SEL1L and HRD1 were identified from the overlapping protein hits passed the selection criteria in SEL1L- and HRD1-immunoprecipitants.

The topology of the interacting proteins in association with the SEL1L-HRD1 complex was mapped based on their dependency on either SEL1L or HRD1. The hits with reduced PSMs in *SEL1L*$^{-/-}$ relative to WT in HRD1 IP-MS, and comparable PSM values in WT and *HRD1*$^{-/-}$ cells in SEL1L IP-MS were categorized hits matching this pattern as SEL1L-dependent hits, referred to as Group I. The hits with comparable PSM values in WT and *SEL1L*$^{-/-}$ cells in HRD1 IP-MS and low *HRD1*$^{-/-}$ to WT PSM ratios in SEL1L IP-MS, were classified as HRD1-dependent hits, or Group II

#### Statistical analysis

Statistics tests were performed in GraphPad Prism version 8.0 (GraphPad Software). Unless indicated otherwise, values are presented as mean ± standard error of the mean (SEM). All experiments have been repeated at least two to three times and/or performed with multiple independent biological samples from which representative data are shown. Statistical differences between the groups were compared using the unpaired two-tailed Student's *t*-test for two groups or one-way ANOVA or two-way ANOVA for multiple groups. $P < 0.05$ was considered statistically significant.

### SUPPLEMENTAL FIGURES AND LEGENDS

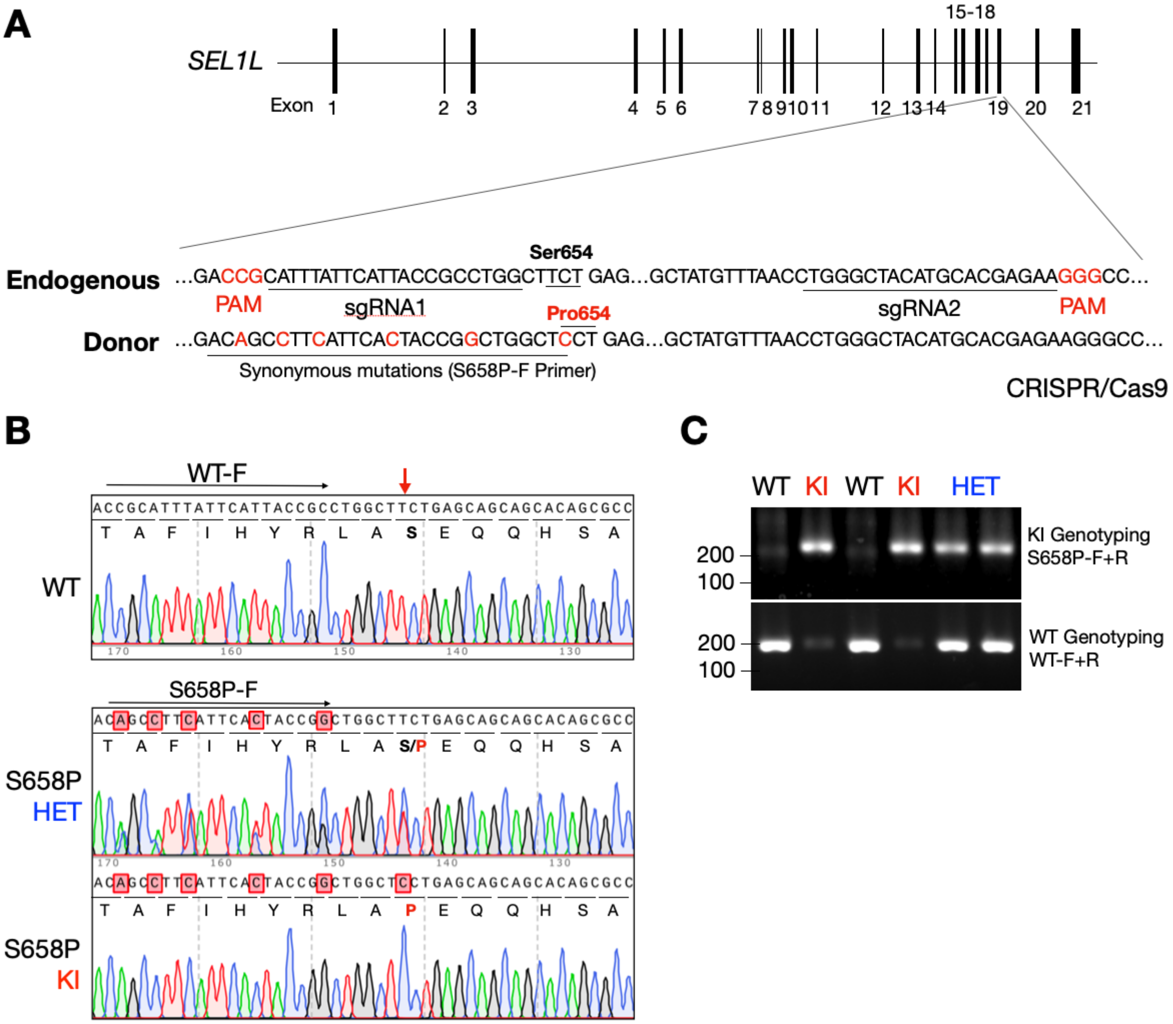


**D**

| Embryonic days (E) | Total | WT | HET | KI |
|---|---|---|---|---|
| Expected ratio | | 25% | 50% | 25% |
| E10.5-E12.5 | 129 | 32 (24.8%) | 75 (58.1%) | 22 (17.1%) |
| E14.5-E16.5 | 116 | 33 (28.4%) | 67 (57.8%) | 16 (13.8%) |
| Post-natal day 21 | 247 | 75 (30.4%) | 145 (58.7%) | 27 (10.9%) |

**Supplemental Figure 1. Generation of *SEL1L*$^{S658P}$ KI mice, related to Figure 1.**
**(A)** Schematic illustration of mouse *Sel1L* gene structures, target loci and donor DNA template sequences. SEL1L S654 in mice is homologous to human S658 (for simplicity, referred to as *SEL1L*$^{S658P}$ for mice throughout the paper). **(B)** Sequence chromatograms for the target site with nucleotide change highlighted in red arrow, and synonymous mutations highlighted in red shaded box. **(C)** PCR genotyping of genomic DNA extracted from P10 mice from two founders. **(D)** Timed pregnancy and frequency of different genotypes showing partial embryonic lethality for the KI mice.

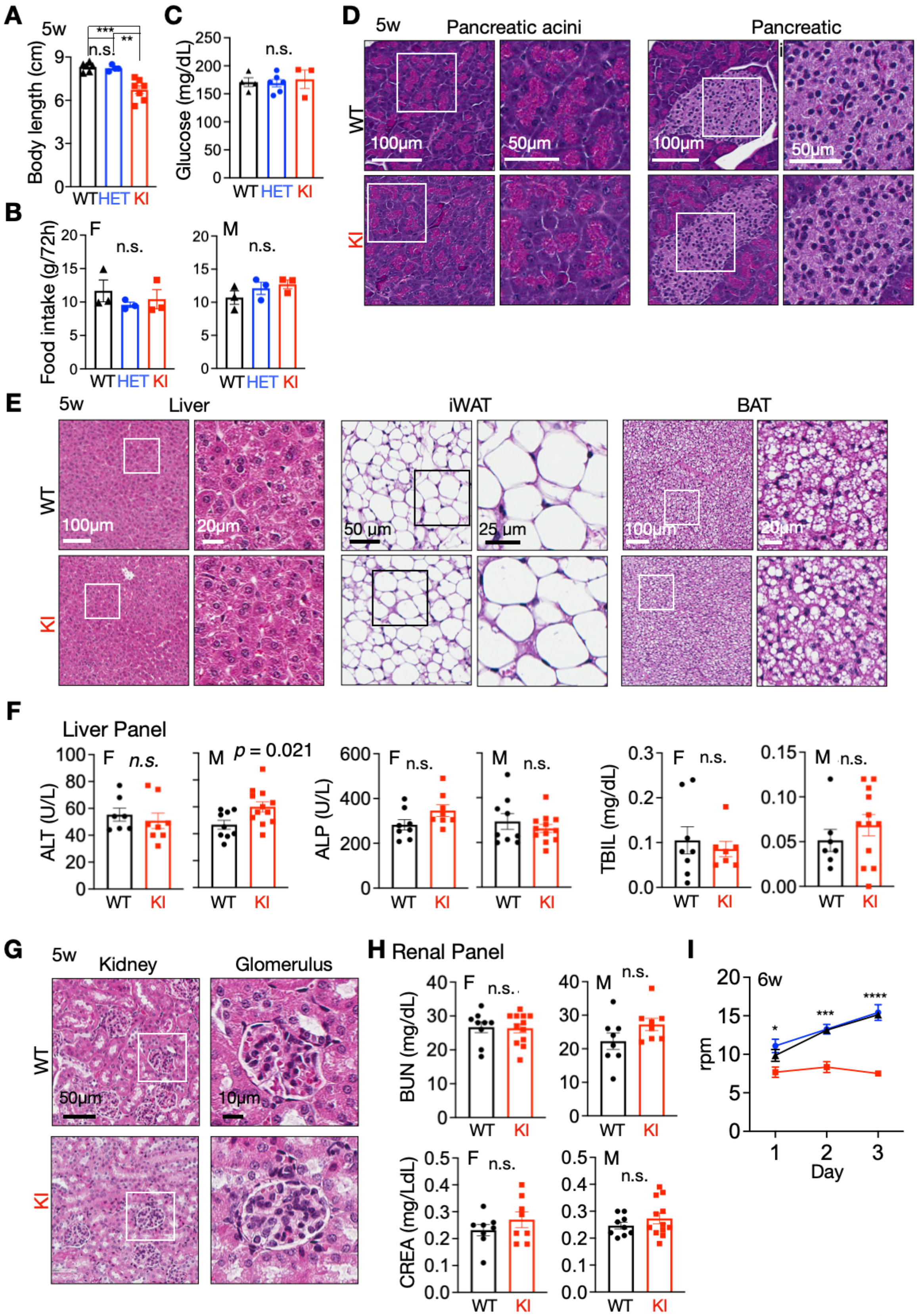


**Supplemental Figure 2. Histological analyses and serum chemistry tests of *SEL1L*$^{S658P}$ KI mice, showing largely normal peripheral tissues, related to Figure 1.**

**(A)** Body length of 5-week-old mice of both genders (n = 4-7 mice per group). **(B)** 72 hr food intake of 5-week-old mice of both genders (n = 3 mice per group). **(C)** Ad libitum blood glucose levels of 5-week-old mice (n = 3-6 mice per group). **(D-E)** Hematoxylin & eosin (H&E) images of various peripheral tissues (**D**, pancreatic acini and islet; **E**, Liver, iWAT and BAT) from 5-week-old mice (n = 3 mice per group). iWAT, inguinal white adipose tissue; BAT, brown adipose tissue. **(F)** Serum levels of alanine transaminase (ALT), total bilirubin (TBIL) and alkaline phosphatase (ALP) in 5-week-old mice of both sexes (n = 7-12 mice per group). **(G)** Hematoxylin & eosin (H&E) stained sections of kidneys and glomerulus from 5-week-old mice (n = 3 mice per group). **(H)** Serum levels of blood urea nitrogen (BUN) and creatinine (CREA) in 5-week-old mice of both sexes (n = 8-12 mice per group). **(I)** Quantitation of rotarod test from mice at 6 weeks of age with 3 days training (n = 7, 12 and 6 mice for WT, HET and KI). Values, mean ± SEM. n.s., not significant; *$p<0.05$, **$p<0.01$, ***$p<0.001$ and ****$p<0.0001$ by one-way ANOVA followed by Tukey's post hoc test (A, B, C), two-tailed Student's *t*-test (F, H) and two-way ANOVA followed by Tukey's multiple comparisons test (I).

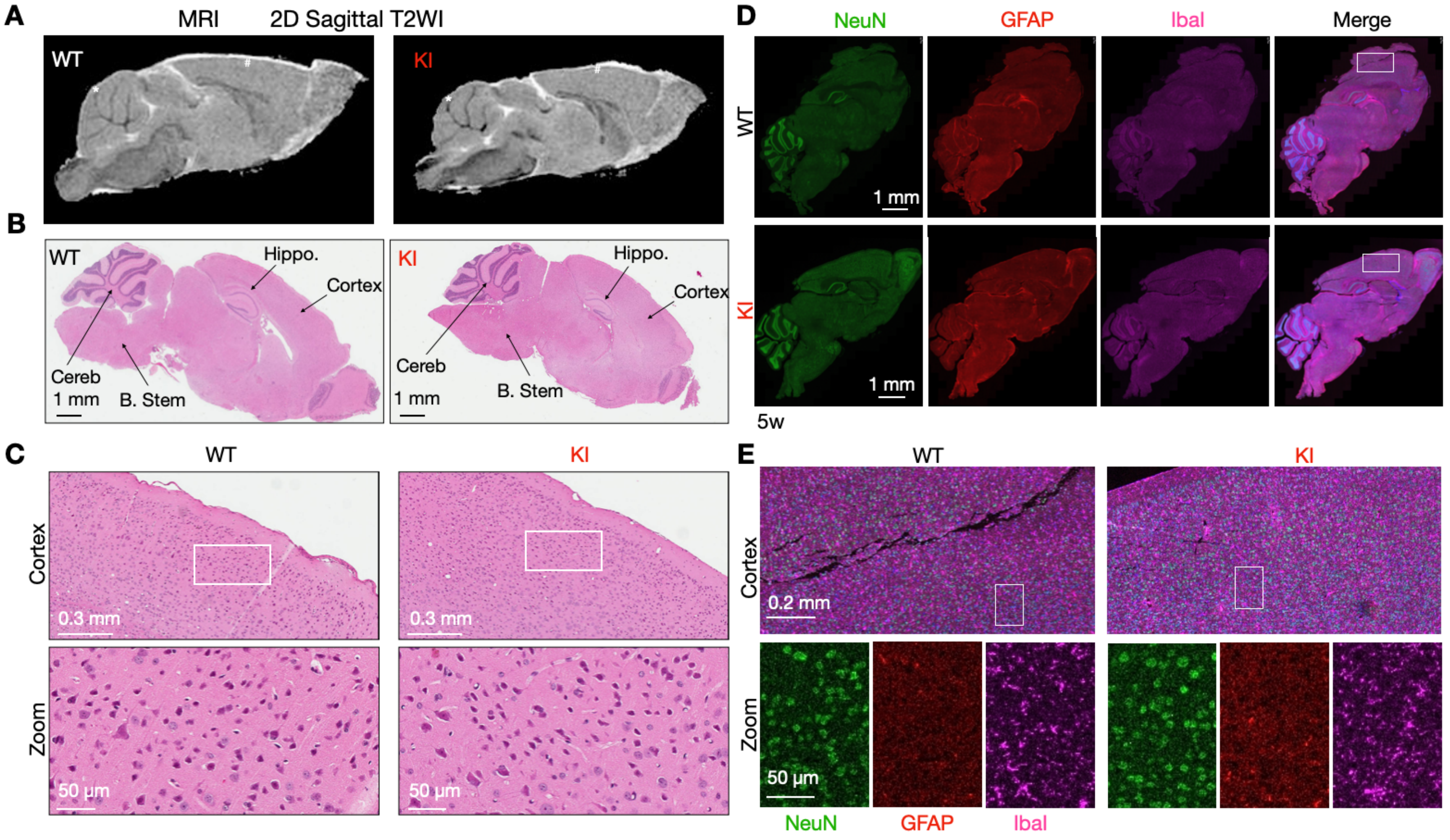


**Supplemental Figure 3. Brains of *SEL1L*$^{S658P}$ KI mice appear largely normal, related to Figure 2.**
**(A)** Morphological sagittal T2-weighted images (T2WI) of 5-week-old mouse brains using MRI analysis (n = 5 mice per group). **(B-C)** Hematoxylin & eosin (H&E) stained sagittal sections of whole brain (**B**) and cortex (**C**) from 5-week-old mice (n = 3 mice per group). Hippo., hippocampus; B. Stem, brain stem; Cereb., cerebellum. **(D-E)** Representative confocal images of NeuN (green), GFAP (red) and Iba1 (purple) staining for neurons, astrocytes and microglia, respectively, in the brains of 5-week-old mice (n = 2 mice per group). Boxed areas are shown in (E).

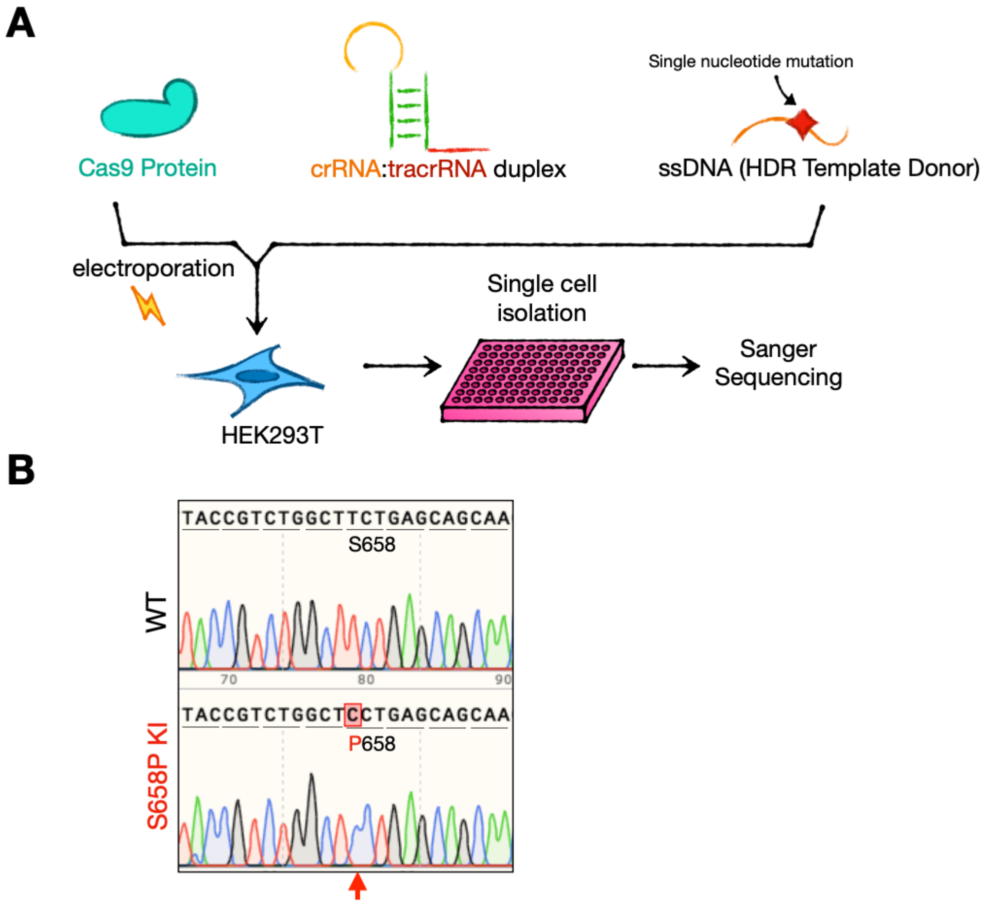


**Supplemental Figure 4. Generation of *SEL1L*$^{S658P}$ knock-in (KI) HEK293T cells, related to Figure 4.**
**(A)** Schematic diagram of the generation of *SEL1L*$^{S658P}$ KI HEK293T using the CRISPR/Cas9 technology and **(B)** Sanger sequencing. The shaded red box and arrow indicated the mutation.

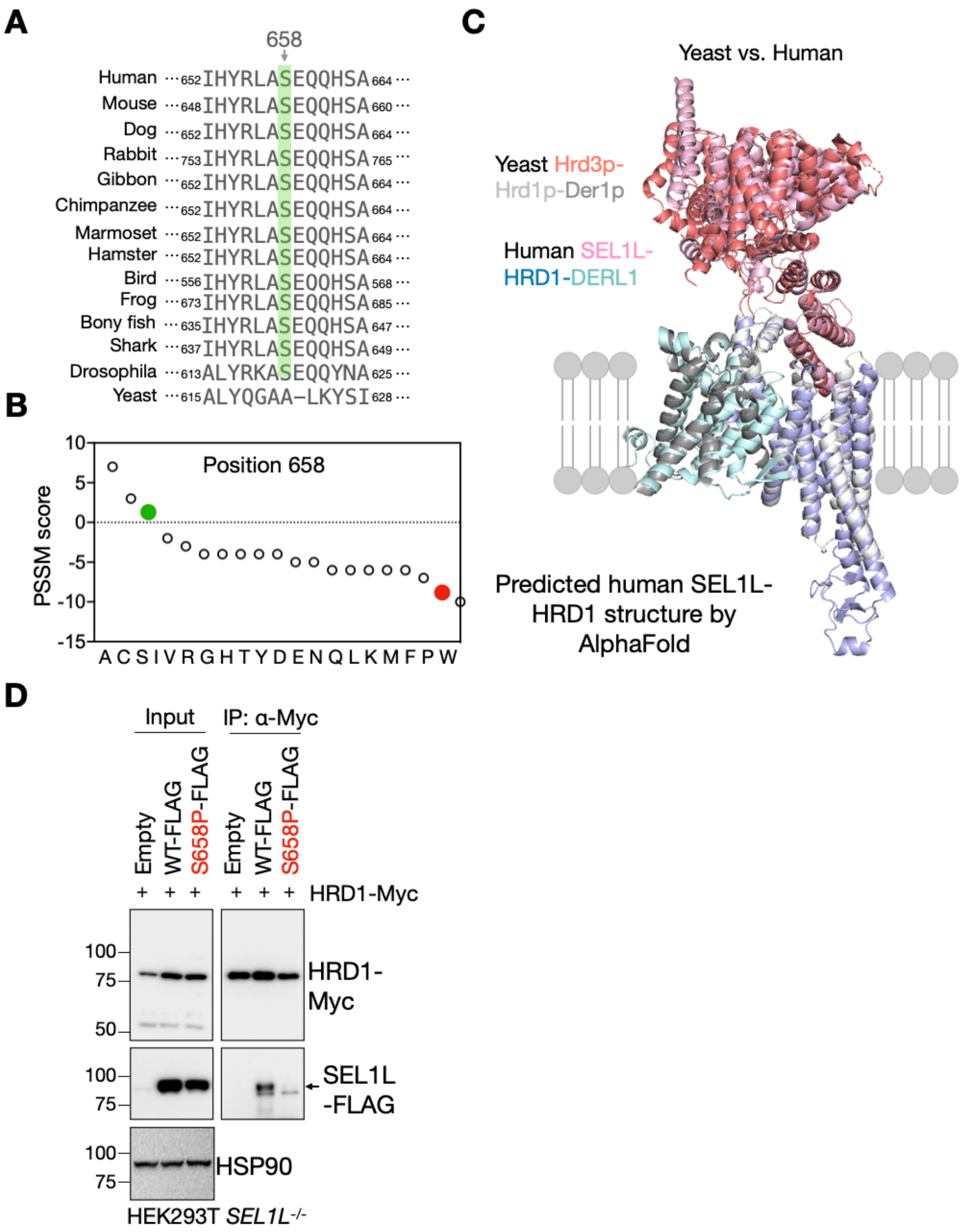


**Supplemental Figure 5. Sequence and structural alignments of SEL1L$^{S658P}$ and the SEL1L-HRD1 complex, related to Figure 5.**
**(A)** Amino acid sequence alignment of SEL1L showing the conservation of SEL1L S658 residue across species (highlighted in green). **(B)** Position-specific scoring matrix (PSSM) score for amino acid of SEL1L 658 position, with S658 in green and P658 in red. **(C)** Structure alignment of the SEL1L-HRD1 complex between human and yeast. **(D)** Immunoprecipitation of Myc-agarose in *SEL1L*$^{-/-}$ HEK293T cells transfected with indicated SEL1L-FLAG and HRD1-Myc constructs to examine HRD1 interaction with SEL1L-FLAG (two independent repeats).

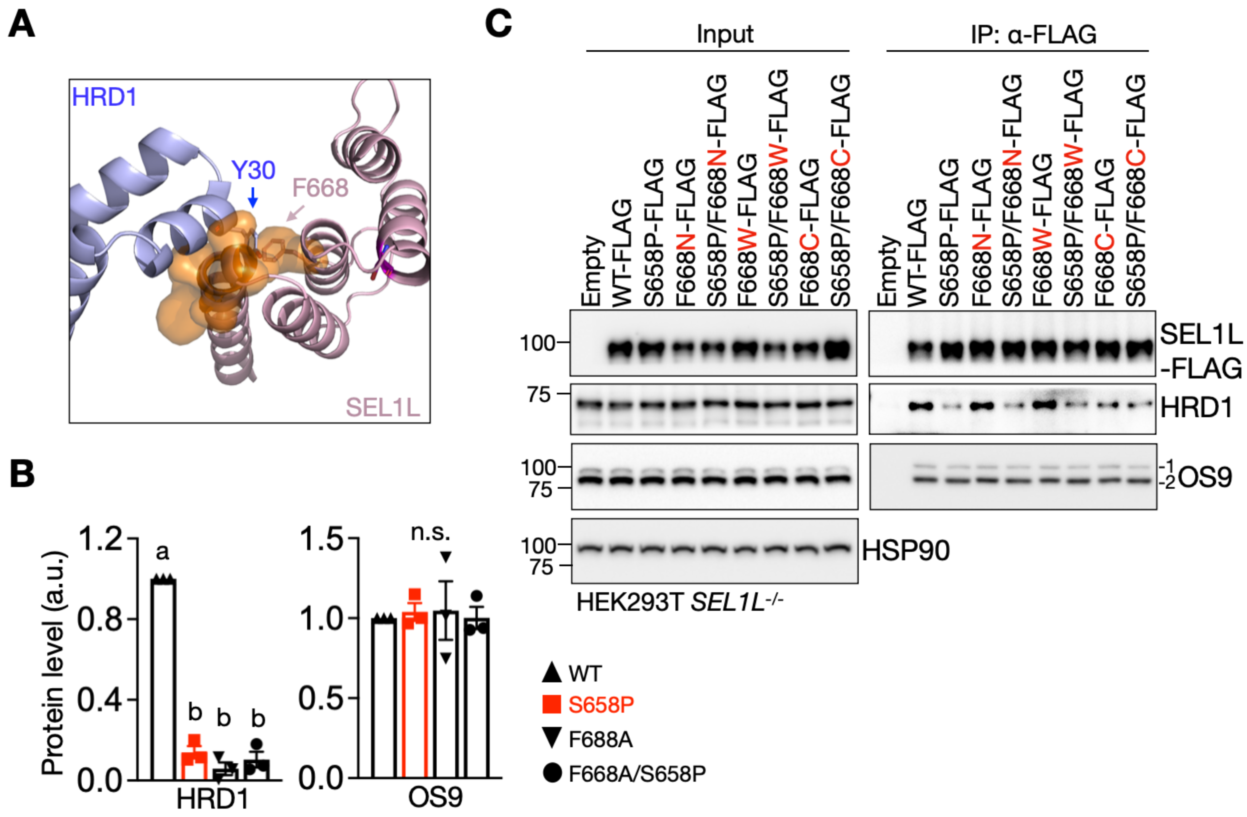


**Supplemental figure 6. Structural and biochemical analyses of SEL1L-F668, related to Figure 6.**
**(A)** side view of SEL1L-HRD1 interaction interface to show HRD1-Y30 and SEL1L-F688 are within the interface. **(B)** Quantitation of HRD1 and OS9 from Figure 5D (n = 3 independent samples per genotype). **(C)** Immunoprecipitation of FLAG-agarose in *SEL1L*$^{-/-}$ HEK293T cells transfected with indicated SEL1L-FLAG constructs to examine the interactions with HRD1 and OS9 (two independent repeats). Values, mean ± SEM. n.s., not significant; Different letters indicate significant differences at the p=0.05 level by one-way ANOVA followed by Tukey's post hoc test (B).

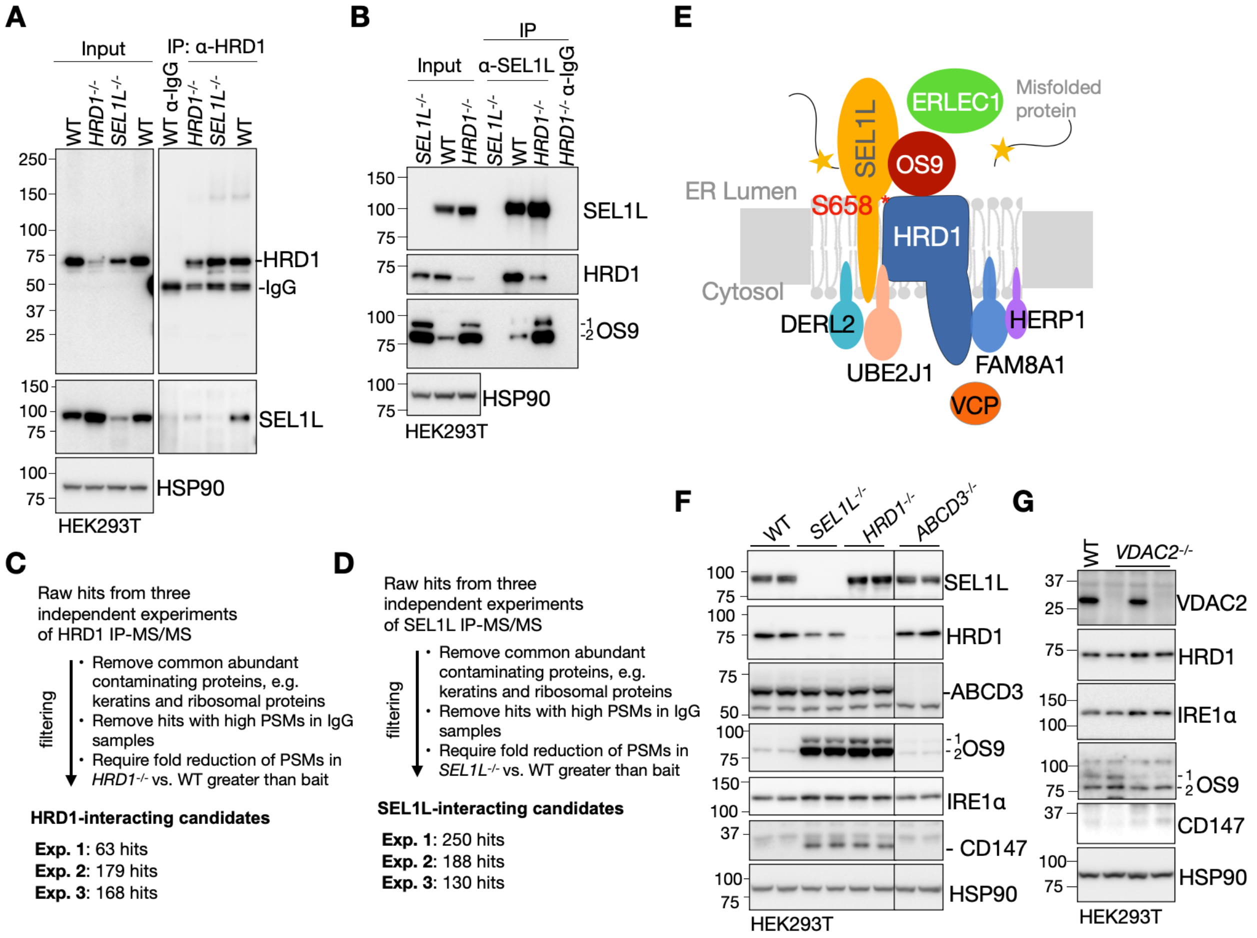


**Supplemental figure 7. The validation and filtering criteria for the IP-MS experiments, related to Figure 7.**

**(A-B)** Immunoprecipitation of HRD1 (A) and SEL1L (B) in WT, *HRD1*$^{-/-}$ and *SEL1L*$^{-/-}$ HEK293T cells to validate the HRD1 antibody or home-made SEL1L antibody in IP. **(C-D)** The flow of the HRD1- (C) and SEL1L- (D) IP-MS strategy to screen for the HRD1/SEL1L-interacting proteins. **(E)** Diagram for the protein components of the SEL1L-HRD1 ERAD complex. **(F-G)** Western blot analysis of indicated proteins in WT, *SEL1L*$^{-/-}$, *HRD1*$^{-/-}$, *ABCD3*$^{-/-}$ and *VDAC2*$^{-/-}$ HEK293T cells showing no effect of VDAC2 and ABCD3 in HRD1 ERAD (two independent repeats). Two panels in (F) were from the same experiment with the irrelevant lanes in the middle cut off.

**Video S1 (separate file): Early-onset ataxia in *SEL1L$^{S658P}$* KI mice.** Balance beam test of 6-week-old mice for WT and KI mice. Animals were trained for two days before recording the test. Loss of coordination and balance in KI mouse is observed at normal speed and after the reduction of four times the normal velocity (slow motion).

**Table S1 (separate file): HRD1-IP mass spectrometry result.** 24 hits for HRD1 interacting proteins from three independent repeats.

**Table S2 (separate file): SEL1L-IP mass spectrometry result.** 47 hits for SEL1L interacting proteins from three independent repeats.
